# Social state gates vision using three circuit mechanisms in *Drosophila*

**DOI:** 10.1101/2024.03.15.585289

**Authors:** Catherine E. Schretter, Tom Hindmarsh Sten, Nathan Klapoetke, Mei Shao, Aljoscha Nern, Marisa Dreher, Daniel Bushey, Alice A. Robie, Adam L. Taylor, Kristin M. Branson, Adriane Otopalik, Vanessa Ruta, Gerald M. Rubin

## Abstract

Animals are often bombarded with visual information and must prioritize specific visual features based on their current needs. The neuronal circuits that detect and relay visual features have been well-studied. Yet, much less is known about how an animal adjusts its visual attention as its goals or environmental conditions change. During social behaviors, flies need to focus on nearby flies. Here, we study how the flow of visual information is altered when female *Drosophila* enter an aggressive state. From the connectome, we identified three state-dependent circuit motifs poised to selectively amplify the response of an aggressive female to fly-sized visual objects: convergence of excitatory inputs from neurons conveying select visual features and internal state; dendritic disinhibition of select visual feature detectors; and a switch that toggles between two visual feature detectors. Using cell-type-specific genetic tools, together with behavioral and neurophysiological analyses, we show that each of these circuit motifs function during female aggression. We reveal that features of this same switch operate in males during courtship pursuit, suggesting that disparate social behaviors may share circuit mechanisms. Our work provides a compelling example of using the connectome to infer circuit mechanisms that underlie dynamic processing of sensory signals.

## Introduction

Behavioral context is critical to how animals detect and interpret visual information. For example, when driving on a congested highway, it is imperative to focus on the movement of the car ahead while ignoring other environmental cues. Such focus or ‘attention’ also occurs during certain behavioral states, including aggression when it is important to be attuned to the movement of a competitor. Pioneering work across primates, rodents, and invertebrates has shown the importance of neuronal populations tuned to specific features, including size, speed, and color (1–11). Recordings in rodents suggest that such visual processing regions receive input from non-sensory areas and behavioral state-dependent neuromodulation (12–17). However, the exact circuit architecture underlying state-dependent gating of visual attention remains unknown.

The fruit fly, *Drosophila melanogaster*, provides a powerful model for the mechanistic dissection of state-dependent visual processing due to its genetic accessibility, brain-wide connectome, and complex behaviors. In flies, visual projection neurons (VPNs) compute the presence and general location of distinct visual features, such as looming or translating objects of varying sizes and speeds (18–20) and relay this information from the optic lobe to different target regions of the central brain (18, 21–23). These visual pathways appear to be highly stereotyped across individuals of both sexes (18, 24). Selective activation of some VPNs gives rise to robust behavioral outputs (18), some of which are context-dependent (25–29). For example, the same looming-responsive VPNs are involved in both landing and takeoff, with the resultant behavioral output determined by octopaminergic modulation of downstream neurons (20, 30, 31).

During social behaviors, detection of conspecifics is critical (32–35). Lobula columnar (LC) neurons, a major class of VPNs, project from the optic lobe to discrete brain glomeruli (18). Three LC cell types, LC9, LC10a, and LC11, that are tuned to fly-sized moving objects, have been implicated in locomotor pursuit of conspecifics (18, 22, 23, 36–42). LC10a is one of four major subtypes of LC10 neurons that each receive distinct inputs in the optic lobe. Previous work has shown that the gain of LC10a responses to fly-size objects increases dramatically during male courtship pursuit (41). Other LC10 subtypes do not appear to display this gain enhancement and their role in social pursuit is less well understood. Stimulation of courtship and aggression promoting P1 neurons (43) in males increases LC10a sensitivity to fly-sized objects, suggesting arousal-dependent modulation of visual processing (41). Yet, in the absence of a male brain-wide circuit diagram, the circuit mechanisms underlying P1 modulation of LC10a remain unresolved. While P1 is only found in males, it represents a subset of the pC1 lineage that gives rise to pC1d and pC1e neurons in females (44–46), which also modulate social states. Recent work has shown that either simultaneous activation of pC1d and pC1e, or activation of the approximately twelve cells comprising the aIPg cell type can generate both acute aggressive behavior and a persistent aggressive state in females (47–49). Our previous work revealed that aIPg provides excitatory input to several LC10 targets, suggesting a role for aIPg in gating the flow of visual information (47). Here, we show that aIPg dedicates a large portion of its synaptic output to modulating visual processing via three circuit mechanisms that regulate multiple visual pathways to facilitate social behaviors (**Figure 1a**).

**Fig. 1.**
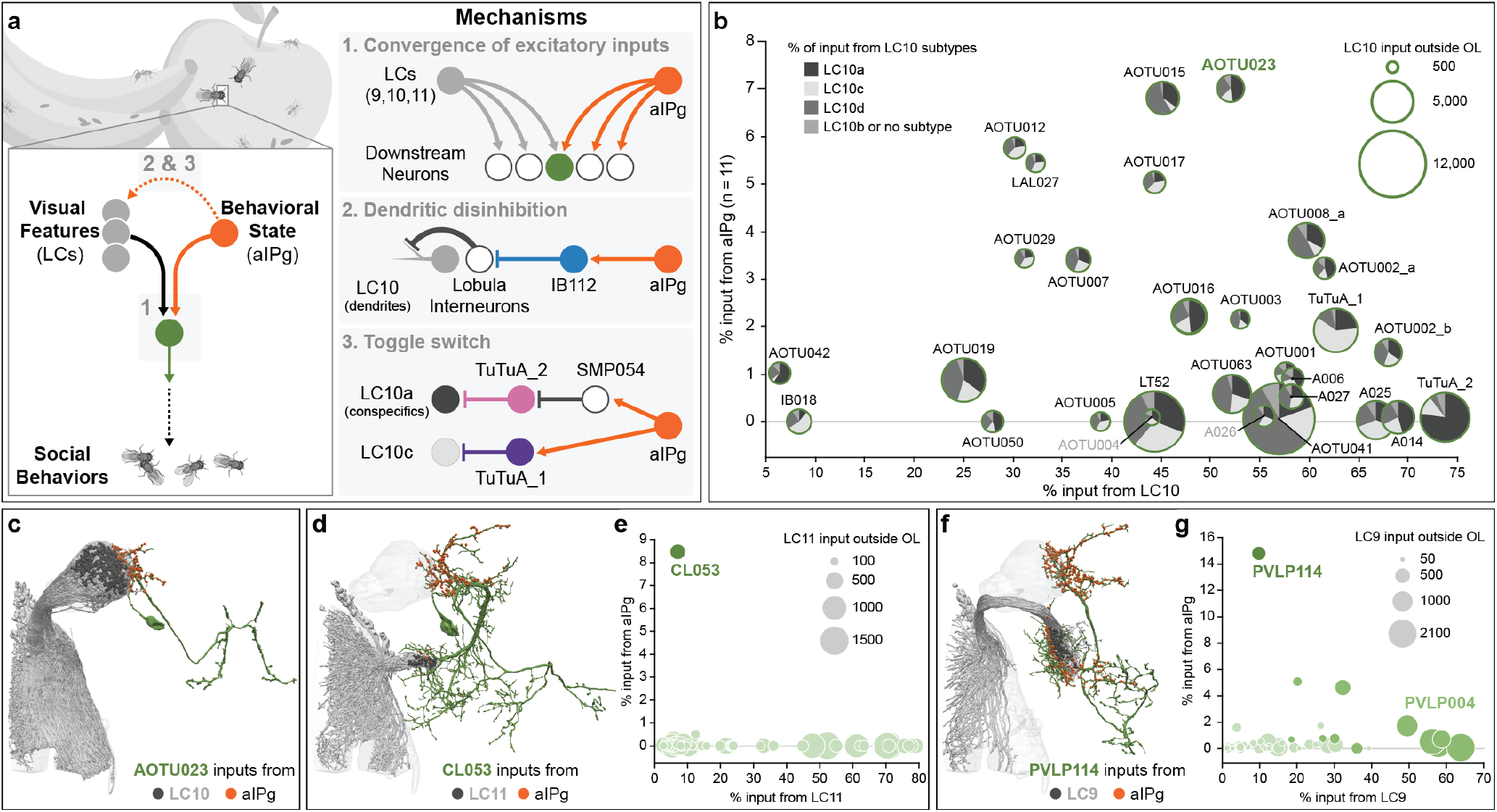
The female aggressive state modifies the flow of visual information by three distinct mechanisms. (a) Summary of the circuit mechanisms that we propose aIPg uses to modulate transmission of visual information by visual projection neurons, specifically LC9, LC10, and LC11. These proposed mechanisms include providing additional excitatory input to a select subset of the direct targets of LC9, LC10, and LC11 (Mechanism 1, Convergence of excitatory inputs), relieving inhibition that acts on LC10 dendritic arbors in the lobula (Mechanism 2, Dendritic disinhibition), and simultaneously flipping a pair of switches that act on the axonal terminals of the LC10a and LC10c cell types to influence which of these two subtypes is active in signaling to downstream targets (Mechanism 3, Toggle switch). Other than the LC9, LC10, and LC11 targets discussed above, only three other neurons get both 1.5% or more of their input from aIPg and 5% or more of their input from an LC type. Arrows indicate putative excitatory connections (cholinergic) and bar endings indicate putative inhibitory connections (GABAergic or glutamatergic). (b) Common shared downstream targets of both aIPg and LC10 neurons. Each target cell type is represented by a circle whose diameter represents the total number of LC10 input synapses it receives. The proportion of those inputs coming from the LC10a, LC10c, LC10d, and other subtypes are indicated as a pie chart. N numbers on axes are per hemisphere. (c) Postsynaptic sites from aIPg (orange) and LC10 (dark gray) on the neuronal outline of AOTU023 (dark green). (d) A diagram of the morphology of CL053 (dark green) is shown with the position of input synapses from aIPg (orange) and LC11 (dark gray). Some ventral arbors lie outside the hemibrain volume and are not shown. (e) Common shared downstream targets of both aIPg and LC11 neurons outside of the optic lobe (OL). Each target cell is represented by a light green circle whose diameter indicates the total number of LC11 synapses that cell receives and whose position on the y-axis represents the percentage of its inputs coming from aIPg and on the x-axis the percentage coming from LC11. This graph shows LC11’s top 51 targets outside the OL representing 74% of its synapses to other cell types outside the OL. (f) A diagram of the morphology of PVLP114 (dark green) is shown with the position of input synapses from aIPg (orange) and LC9 (dark gray). Some ventral arbors lie outside the hemibrain volume and are not shown. (g) Common shared downstream targets of aIPg and LC9 neurons outside of the OL. Each target cell is represented by a light green circle. The diameter of each circle indicates the total number of LC9 synapses that cell receives and whose position on the y-axis represents the percentage of its inputs coming from aIPg and on the x-axis the percentage coming from LC9. This graph shows LC9’s top 54 targets outside the OL representing 83% of its synapses to other cell types outside the OL.

## Results

### Vision is critical for aggression

Multisensory cues are important for locating others and directing aggressive actions (34, 47). Previous behavioral evidence suggested the importance of visual information in aIPg-induced aggression (47). We confirmed this by eliminating the ability of females to receive visual information using a mutation in the norpA gene which encodes a key component of the phototransduction pathway (50). Activating aIPg in norpA mutant females did not result in continued aggressive interactions, even after they made physical contact (**Supplementary Figure 1a – c**). These data emphasize the importance of visual cues in aggressive interactions elicited by aIPg activation.

### Shared targets of vision and internal state

To determine which visual pathways are modulated by aIPg, we performed a comprehensive analysis of the female connectome (24) and identified neurons receiving input of over 100 synapses from both aIPg and visual projection neurons (VPNs). We found that the VPNs participating in this circuit motif were a small subset (8/44) of lobula columnar (LC) cell types. Moreover, each of these eight LC cell types is known to be responsive to small moving and/or looming objects (18, 22, 23). This combination of aIPg and LC inputs may endow the downstream neurons with the capacity to integrate an aggressive internal state and socially relevant visual information (**Figure 1a**). LC10 shares the largest number of neuronal targets with aIPg. We identified 23 cell types that receive more than 25% of their input synapses from LC10 neurons and about half of these also receive more than 2% of their input synapses from aIPg (**Figure 1b**). Each of these shared outputs of aIPg and LC10 receives input from all LC10 subtypes, although in different proportions (**Figure 1b – c**). Aside from LC10, aIPg shares primarily one downstream target with LC11 (CL053) and two downstream targets with LC9 (PVLP114 and PVLP004) (**Figure 1d – g**). Only three other downstream neurons receive significant input from both LC and aIPg neurons: PVLP120 receives 30% of its synaptic inputs from LC17, 19% from LC12 and 1.5% from aIPg; SMP312 receives 5.3% of its inputs from LC21 and 4.6% from aIPg; and PVLP006 receives 35% of its inputs from LC6, 11% from LC16 and 2.3% from aIPg. Thus, these downstream targets of both LCs and aIPg, have the capacity to receive both excitatory visual and aIPg input during an aggressive encounter.

The proportions of input from aIPg and the LCs vary (**Figures 1b, e, g**). The vast majority of cell types that receive LC10 input are interneurons in the AOTu, which then connect to descending pathways that drive motor action (38, 51, 52). These interneurons therefore may control distinct facets of aggressive behavior. Genetic reagents that allow us to target AOTu cell types, and perhaps combinations of cell types, as well as assays for subtle aspects of behavior will be needed to further explore these parallel pathways.

Previous work characterizing visual responses of different LC populations using in vivo calcium imaging revealed LC11 to be selectively tuned to small moving objects (approximately 4.5° in angular width and height as subtended on the retina) (22, 23, 42). For comparison, we used the same experimental approach to examine visual feature selectivity of LC9 and LC10a. Similar to LC11, LC9 preferred smaller moving objects (approximately 4.5° in width and approximately 2° in height) than LC10a (approximately 15 – 30° in width and height) (**Figure 2a and Supplementary Figure 2a – b**). As perception of an object’s size depends on its distance, such variations in size selectivity suggest differential activation of LC9 and LC10a when a female is close to versus far away from another fly. Additionally, LC9 displayed a prolonged calcium response to slow dark looming stimuli (**Supplementary Figure 2c**). Taken together, our results show that LC9, LC10a, and LC11 are among the limited subset of LC neurons tuned to fly-sized moving objects and suggest that these LCs may be differentially activated over time, as a function of varying inter-fly distance, during aggressive encounters.

**Fig. 2.**
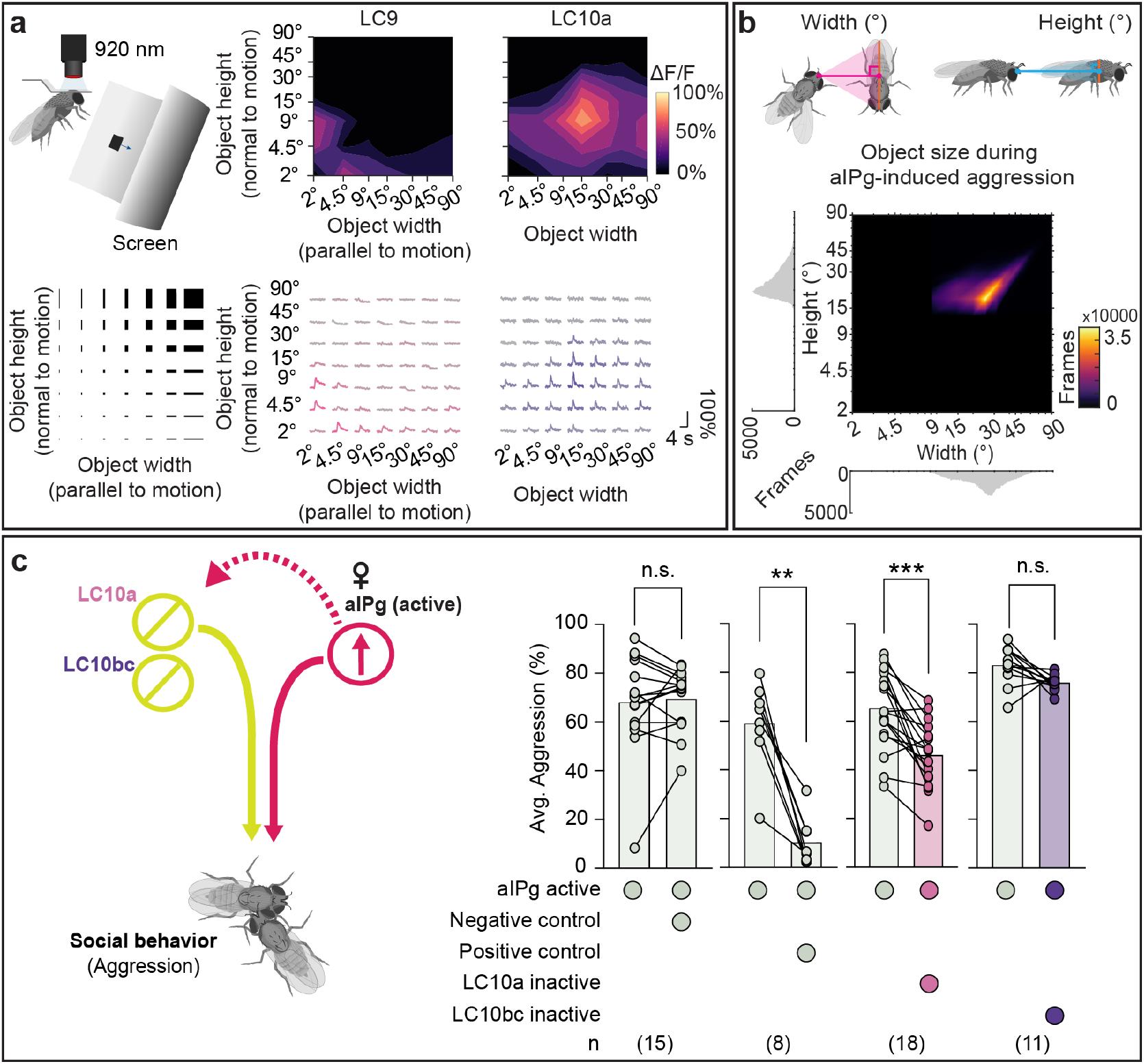
LC10a is tuned to medium-sized moving objects, similar to those found during female aggression. (a) Schematic of experimental setup (top left) for presentation of moving dark rectangles of parameterized spatial dimensions (bottom left). Receptive-field centers were mapped for individual axons leaving the lobula, and each stimulus was then translated across the entire receptive field (see **Supplementary Figure 2a – b**). Average traces for individual LC9 axons in females and LC10a axons in males (bottom right). Average heat map representations of peak responses are shown across multiple animals (top right). LC9: n = 4 flies, n = 4 neurons. LC10a: n = 5 flies, n = 7 neurons. (b) Heat map representations of conspecific angular sizes experienced during aIPg-induced female aggression. During female aggression, the mean conspecific size as subtended on the retina was 26.1 +/- 8.8° (mean +/- standard deviation) in height and 32.1 +/- 13.0° in width. Female aggression frames were defined using the JAABA aggression classifier and calculated from 79 trajectories. Illustrations on top depict calculations for angular width and height of target female as subtended on subject female’s retina. See **Supplementary Figure 2f** for angular position and velocity data. (c) Average time spent performing aggressive behaviors before and during stimulus periods in which a 30 s continuous green (9 mW/cm^2^) light stimulus was delivered. See **Supplementary Figure 2g - i** for for time course and non-permissive temperature controls. The following genotypes were used: aIPg-LexA > TrpA emptySS > GtACR (aIPg active Negative control), aIPg-LexA > TrpA aIPg-SS > GtACR (aIPg active Positive control), aIPg-LexA > TrpA LC10a-SS > GtACR (aIPg active LC10a inactive), and aIPg-LexA > TrpA LC10bc-SS1 > GtACR (aIPg active LC10bc inactive). The average for the pre-stimulus period was calculated using the first (last 15 s) pre-stimulus period based on the time course data (see **Supplementary Figure 2h – i**). Averages were calculated over all flies in an experiment, with each dot representing one experiment containing approximately seven flies. All data points are shown to indicating the range and top edge of bar represents the mean. In the diagram on the left, cell types inactivated with GtACR are circled in yellow and those activated with TrpA are circled in red. Data were pooled from four independent replicates, which included separate parental crosses and were collected on different days. A non-parametric Wilcoxon Matched-pairs Signed Rank test was used for statistical analysis. Asterisk indicates significance from 0: **p<0.01; ***p<0.001.

LC10a has been implicated in male courtship behavior (40, 41), yet LC10a’s role in female aggression has not been explored. To explore this potential role for LC10a, we first examined the visual object sizes and speeds experienced by female flies during aggressive encounters. During periods of aIPg-mediated aggression, female flies modulate their velocity with respect to another fly such that the angular size of the nearest fly remains 32.1 +/- 13.0° in width (mean +/- standard deviation) and 26.1 +/- 8.8° in height, which corresponds to the flies being less than a body length apart (**Figure 2b**). Such stimuli are within the preferred ranges of object size and speed of LC10a neurons (**Figure 2a and Supplementary Figure 2d, f**), consistent with the notion that LC10a could play a role during female aggression. The preferred range of stimuli for LC10a are also within the range of object sizes (approximately 29° in width, approximately 16° in height) courting males fixate on during courtship (**Supplementary Figure 2e - f**), similar to previous work (40). Moreover, optogenetic inactivation of LC10a, but not LC10bc, resulted in a sustained decrease in aIPg-mediated aggressive behaviors, including individual component features such as touch (**Figure 2c and Supplementary Figure 2g – i**). The distance between LC10a-silenced flies and others increased, and the angular size subtended on the retina by the nearest fly correspondingly decreased (**Supplementary Figure 2h**). Collectively, these results suggest that LC10a plays a similar role in the visual tracking of social targets in both male courtship and female aggressive encounters.

### Broad disinhibition via a centrifugal neuron

Our connectomic analyses suggest a second mechanism by which aIPg enhances information flow from LC10 neurons (**Figure 1a**). IB112, aIPg’s second strongest target based on synapse number, has nearly 90% of its synaptic output in the lobula (**Figure 3a**), suggesting a role in modulating visual inputs to the brain. To examine its neurophysiology and behavioral contributions, we generated two cell-type-specific driver lines for IB112 and confirmed this cell type to be glutamatergic, suggesting that it is likely inhibitory (53) (**Supplementary Figure 3a – h**). Whole-cell patch-clamp recordings of IB112 during aIPg stimulation in female brain explants confirmed direct functional connectivity between aIPg and IB112 (**Supplementary Figure 4a**). To test the role of such connections in female aggressive behaviors, we performed epistasis experiments by thermogenetically activating aIPg while optogenetically silencing IB112. Inactivation of IB112 resulted in a prolonged decrease in aggression, including touch and other component behavioral features, during silencing (**Figure 3b and Supplementary Figure 4b – d**), confirming the importance of IB112 outputs in the lobula.

**Fig. 3.**
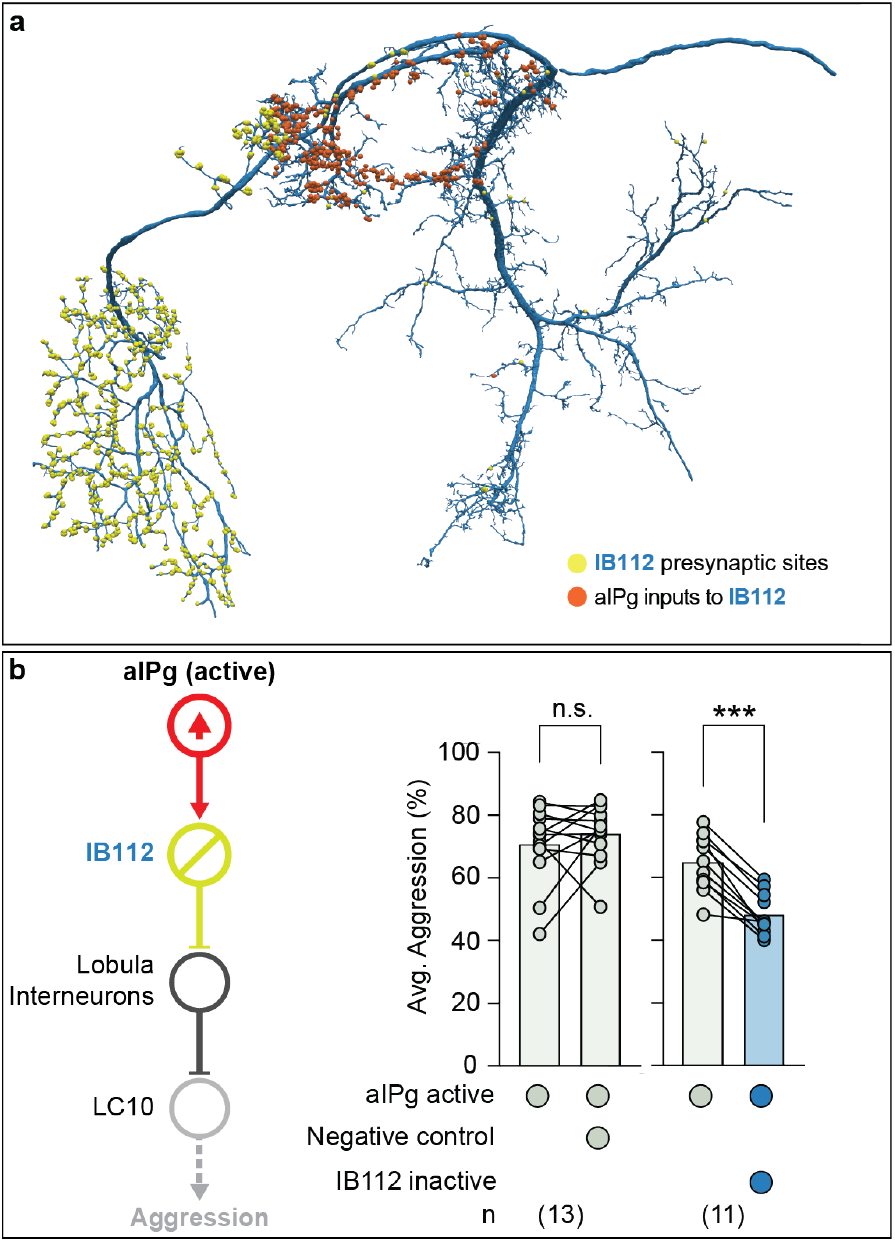
Polysynaptic connections from aIPg to the lobula shape aggressive behaviors. (a) Postsynaptic sites from aIPg (orange) and presynaptic sites of IB112 to its downstream targets (yellow) in the lobula are shown on the neuronal outline of IB112 (dark blue). (b) Average time spent performing aggressive behaviors before and during stimulus periods in which a 30 s continuous green (9 mW/cm^2^) light stimulus was delivered. See **Supplementary Figure 4b – d** for time course and non-permissive temperature controls. Averages were calculated over all flies in an experiment, and each dot represents one experiment containing approximately seven flies. All data points are shown to indicating the range and top edge of bar represents the mean. The following genotypes were used: aIPg-LexA > TrpA emptySS > GtACR (aIPg active Negative control) and aIPg-LexA > TrpA IB112-SS2 > GtACR (aIPg active IB112 inactive) (see **Supplementary Figure 4c - d** for Positive Control and IB112-SS1 data). In the diagram on the left, cell types inactivated with GtACR are circled in yellow and those activated with TrpA are circled in red. IB112 and the relevant lobula interneurons are predicted to be glutamatergic and are presumed inhibitory (see **Supplementary Figure 3h** for confirming data on IB112’s neurotransmitter expression). Data were pooled from four independent replicates, which included separate parental crosses and were collected on different days. A non-parametric Wilcoxon Matched-pairs Signed Rank test was used for statistical analysis. Asterisk indicates significance from 0: ***p<0.001.

We used the recently completed optic lobe connectome in males (54) to examine the synaptic outputs of IB112 in this dataset (**Figure 6 and Supplementary Figure 9**) and then verified the presence of these connections in the less extensively annotated female optic lobe connectome (55). Within the male and female lobula, IB112 forms strong connections with three neurons predicted to be glutamatergic: Li22, Li14, and LT52 implying these connections are likely to be inhibitory (53). The lobula local interneuron Li22 was the most predominant IB112 target, receiving nearly 20% of all IB112 output synapses. These IB112 synapses make up 86% of synaptic input to Li22 in males, and 80% of synaptic input to Li22 in female. IB112 contributes substantial, but markedly less input to Li14 and LT52 in both sexes. In the male, this corresponds to each Li22 cell, on average, getting 30 synapses from IB112 as compared to only 3 connections for Li14 and 13 for LT52. Li22, Li14, and LT52 each then synapses on to the dendrites of all LC10 subtypes (for additional details see **Figure 6**).

Analysis of the connectome has identified more than 70 similar cell types, so-called centrifugal neurons, which have most of their inputs within the central brain and their outputs in the optic lobes (54, 55). While two previous examples of centrifugal neurons have been predicted to use neurotransmitters, such as octopamine, to modify behavior (56, 57), their circuit-level functions are unknown. Thus, IB112 provides the first example of this important class of inputs to the optic lobe where not only a behavioral role but also the relevant direct synaptic inputs in the central brain and outputs in the optic lobe have been identified.

### An axo-axonal-mediated toggle switch

Our previous work identified the TuTuA neurons as potential mediators of aIPg regulation of LC10 signaling in the AOTu (47). Further analysis of the female connectome revealed that there are two subtypes of TuTuA neurons, referred to as TuTuA_1 and TuTuA_2, each represented by a single glutamatergic cell per brain hemisphere. We found that the TuTuA subtypes have distinct patterns of connectivity that suggest they differentially gate LC10a and LC10c (**Figure 4a**). Given the connectivity and predicted neurotransmitters in this circuit, our simplest interpretation is as follows: when active, aIPg provides excitatory input to TuTuA_1, which forms axo-axonal connections with LC10c inhibiting its ability to transmit information (**Figure 4b – c**). In addition, aIPg indirectly targets TuTuA_2 through a GABAergic interneuron, SMP054. TuTuA_2 forms axo-axonal inhibitory connections with LC10a. This disynaptic sequence of inhibitory connections serves to disinhibit LC10a transmission (**Figure 4b – d**). In this simple view, aIPg activation has the potential to oppositely modulate LC10a and LC10c and their respective downstream circuits. Should aIPg have only two states – active and silent – the Tu-TuA neurons would act as a pair of opponent switches that flip from LC10c to LC10a transmission when aIPg becomes active.

**Fig. 4.**
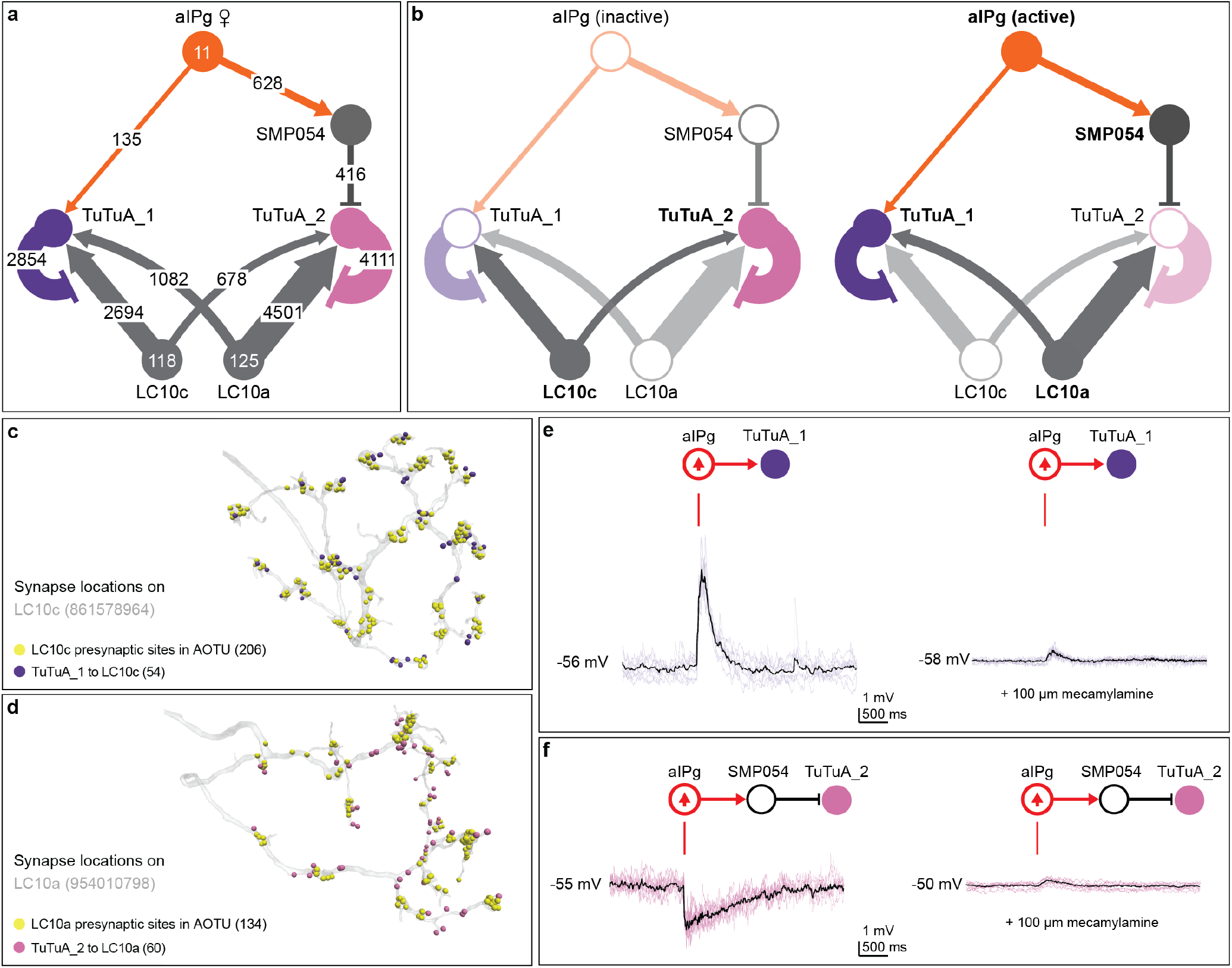
aIPg selectively amplifies LC10a, while dampening LC10c transmission via TuTuA neurons. (a) Connectivity between aIPg, TuTuA subtypes (TuTuA_1, TuTuA_2), LC10a, and LC10c. Exact synapse numbers are indicated on the arrows, which are also scaled in size according to synapse counts. Note that the TuTuA_1 and TuTuA_2 neurons are highly specific in their connections for LC10c and LC10a, respectively: 97% of TuTuA_1’s synapses onto LC10 go to the LC10c subtype, whereas 97% of TuTuA_2’s synapses onto LC10 go to LC10a subtype. Arrows indicate putative excitatory connections (cholinergic) and bar endings indicate putative inhibitory connections (SMP054, GABAergic; TuTuA_1 and TuTuA_2, glutamatergic). (b) Predicted outcomes for circuit dynamics based on aIPg activity. See text for details. Cells and connections with higher predicted activity are displayed in bold and dark colors. (c – d) Axo-axonal synapses between TuTuA_1, TuTuA_2, LC10a, and LC10c on representative neuronal skeletons for LC10c (Body ID: 861578964) and LC10a (Body ID: 954010798). Note how inhibitory synapses from the TuTuA neurons are interspersed with the LC10’s output synapses. (e) Excitatory responses recorded from TuTuA_1 (n = 16 cells) using patch clamp electrophysiology in female brain explants before, during, and following a 2 ms stimulation of aIPg neurons. The excitation was largely abolished by mecamylamine, a n-AchR blocker. Individual trials in purple (n = 8 trials from 1 cells), mean in black. (f) Inhibitory responses recorded from TuTuA_2 (n = 16 cells) before, during, and following a 2 ms stimulation of aIPg neurons. The inhibition was completely removed by mecamylamine. Individual trials are in pink (n = 8 trials from 1 cell), mean is in black. In the diagrams above the traces, cell types activated with CsChrimson are circled in red and those recorded from are in purple or pink depending on the TuTuA subtype.

TuTuA_1 and TuTuA_2 are predicted to have opposite effects on the activity of LC10a and LC10c (**Figure 4b**). Excitatory feedback from LC10a and LC10c to TuTuA_1 and TuTuA_2, respectively, likely reinforces this opponency (**Figure 4b**). Specifically, the connectome revealed that a major downstream target of LC10a is TuTuA_1, the same TuTuA that inhibits LC10c. Thus, when LC10a is active it serves to reinforce the suppression of LC10c activity. Thus, when LC10a is active it serves to reinforce the suppression of LC10c activity. Analogous to LC10a’s connections to TuTuA_1, a major LC10c target is TuTuA_2. This complementary indirect pathway serves to suppress LC10a activity when LC10c is active.

Features of this circuit cannot be fully understood from the connectome alone. For example, both TuTuAs provide axo-axonal inhibition on distinct subsets of LC10 neurons and receive substantive excitatory feedback from those same LC10s. It is conceivable that these serve as feedback inhibition motifs (58) to regulate the gain of TuTuA output (59, 60). Furthermore, how the neurons of this circuit integrate synaptic connections over time and at a subcellular level remain unknown and would require pointed neurophysiological interrogation beyond current technical capacities.

We were able to test many of the predictions from the circuit diagram by generating GAL4 driver lines specific for each TuTuA subtype (**Supplementary Figure 5a – w**), and subsequently using these genetic reagents in functional assays. Electrophysiology and calcium imaging during aIPg stimulation confirmed the following predicted connections: direct aIPg excitatory connections to TuTuA_1; indirect inhibitory connections to TuTuA_2; and excitatory connections of LC10a to both TuTuA_1 and TuTuA_2 (**Figure 4e – f and Supplementary Figure 6a – d**). pC1d and pC1e neurons, previously shown to be upstream of aIPg in the female aggression circuit (47), were also found to provide indirect inputs to TuTuA_2 through SMP054 (**Supplementary Figure 6e – i**).

To test the behavioral effects of this circuit architecture, we acutely silenced TuTuA_1 during aIPg activation. As expected, we found that acute optogenetic inhibition of TuTuA_1 activity during chronic aIPg activation transiently decreased aggression (**Figure 5a and Supplementary Figure 7a, e**). Next, we optogenetically activated TuTuA_2 during chronic aIPg thermogenetic activation and found that it significantly reduced female aggressive behavior during the duration of the stimulus and correspondingly increased the distance between individuals (**Figure 5a and Supplementary Figure 7b – e**). Furthermore, TuTuA_2 optogenetic inactivation increased aggression behavior in the absence of aIPg activation (**Figure 5b and Supplementary Figure 7f**). Taken together, this work provides strong evidence in support of a novel toggle switch mechanism whereby aIPg activation shifts the relative gain of the LC10a and LC10c visual pathways.

**Fig. 5.**
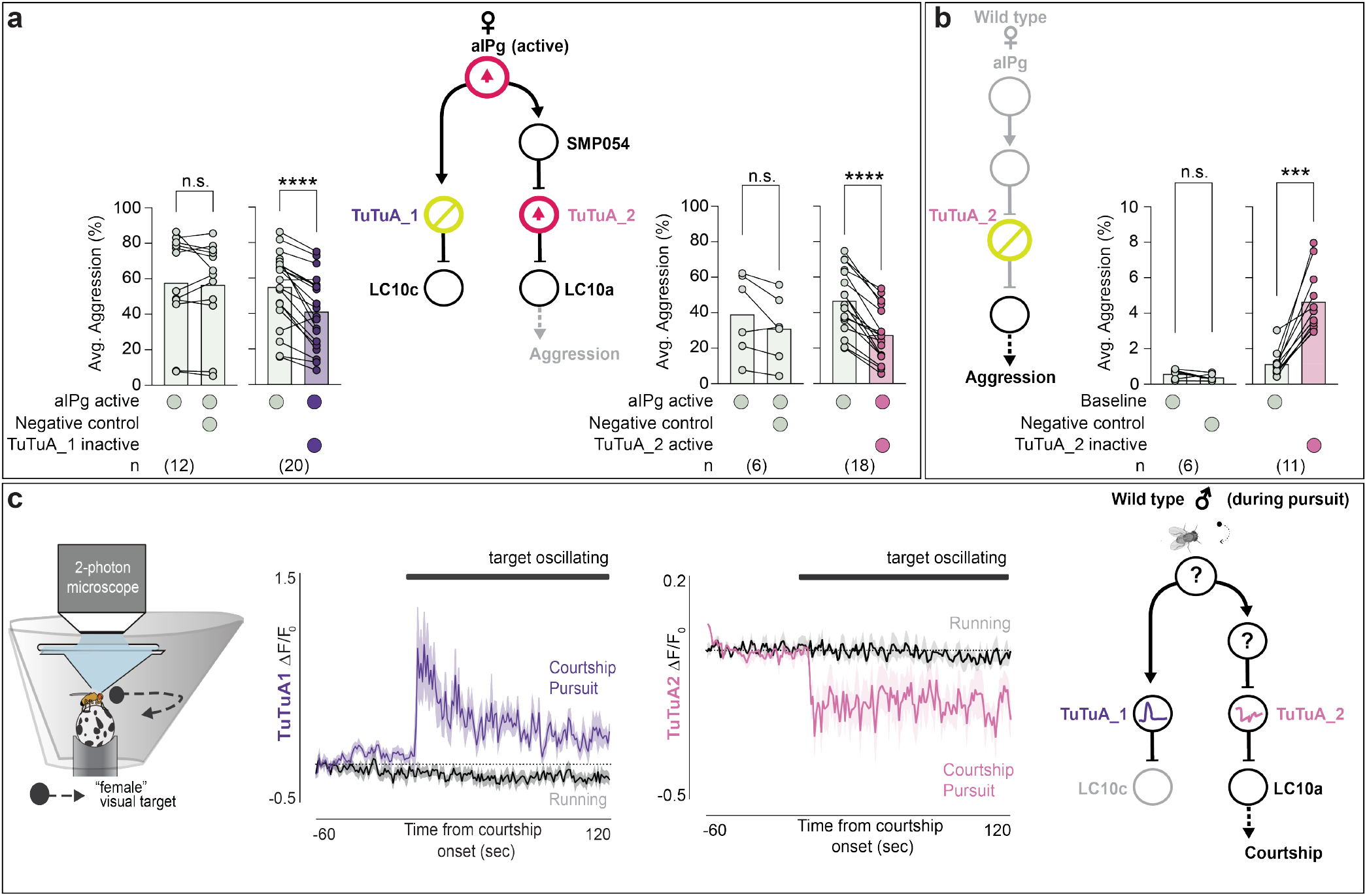
Selective modulation of LC10 subtypes shapes female aggression and male courtship. (a – b) Average time spent performing aggressive behaviors before and during stimulus periods in which a 30 s continuous green (9 mW/cm^2^) or red (3 mW/cm^2^) light stimulus was delivered. See **Supplementary Figure 7a – b, and e** for time course and non-permissive temperature controls. The following genotypes were used: (a, left panel) aIPg-LexA > TrpA emptySS > GtACR (aIPg active Negative control), aIPg-LexA > TrpA TuTuA_1-SS > GtACR (aIPg active TuTuA_1 inactive) (see **Supplementary Figure 7a** for Positive Control); (a, right panel): aIPg-LexA > TrpA emptySS > CsChrimson (aIPg active Negative control), aIPg-LexA > TrpA TuTuA_2-SS > CsChrimson (aIPg active TuTuA_2 active); (b): emptySS > GtACR (Negative control), TuTuA_2-SS > GtACR (TuTuA_2 inactive). Based on the time course data (see **Supplementary Figure 7a – b, f**), either the last 10 s (a, left panel) or 30 s (a, right panel and b) of each pre-stimulus period was compared to averages across all three stimulus periods (first 10 s of each – left panel a, or the 30 s of each – right panel, a and b). Inactivation experiments in (b) were performed with group housed flies which have a decreased baseline in aggression. Averages were calculated over all flies in an experiment, with each dot representing one experiment containing approximately seven flies. All data points are shown to indicating the range and top edge of bar represents the mean. Cell types inactivated with GtACR are circled in yellow and those activated with either TrpA or CsChrimson are circled in red. Data were pooled from four (a, left panel) and two (a, right panel and b) independent replicates, which included separate parental crosses and were collected on different days. (c) Left: schematic of the visual virtual reality preparation for male courtship (redrawn from (41)). Males walking on an air-supported foam ball are presented with a dynamic fly-sized visual target that sweeps left and right across the visual panorama at regular intervals. Center: Responses of TuTuA neurons (average Δ*F* /*F*_0_ ) to a visual target during periods of courtship pursuit (purple or pink) or general locomotion (black). The mean is represented as a solid line and shaded bars represent standard error between experiments (TuTuA_1-SS1, n = 4 flies; TuTuA_2-SS1, n = 5 flies). Black line above indicates when the visual target was oscillating. Courtship is determined by the vigor of male pursuit and the presence of unilateral wing-extensions. Right: The schematic represents circuit activity during male courtship pursuit. Cell types with question marks in the schematic are not definitively known due to the lack of the male connectome. A non-parametric Wilcoxon Matched-pairs Signed Rank test was used for statistical analysis. Asterisk indicates significance from 0: ***p<0.001; ****p<0.0001.

It is interesting to note that the targets of LC10 within the AOTu are regulated in two ways by aIPg (**Figure 1a, Convergence of excitatory inputs and Toggle switch**). First, aIPg provides significant direct input to about half of these AOTu interneurons (**Figure 1b**). Second, aIPg activation is predicted to produce a global shift in the visual input those neurons receive—even those that do not receive direct aIPg inputs—by gating which LC10 subtypes can effectively signal to them (**Figure 4b**). Thus, aIPg is primed to regulate the flow of visual information through all visual AOTu interneurons using either one or both of these distinct circuit mechanisms.

### The same TuTuA-mediated switch functions during male arousal

Previous work has demonstrated that P1 neu-rons, directly or indirectly, increase the gain of LC10a activity, but not LC10c activity, in the AOTu during courtship pursuit (41). We previously suggested that a common mechanism involving the TuTuA neurons might underlie state-dependent gating in females and males (47). Consistent with this suggestion, we show that TuTuA neurons exhibit similar morphology and baseline physiological properties across sexes (**Supplementary Figure 5p – s and Supplementary Figure 8a – b**). We then examined TuTuA activity in tethered males as they spontaneously initiated courtship pursuit of a ‘fictive female’ represented as a high contrast dot that moves at a constant angular velocity across the male’s visual field (41) (**Figure 5c**). While both TuTuA neurons were insensitive to the visual profile of the fictive female when males viewed it passively, the onset of courtship marked a striking change in their calcium activity: the activity of TuTuA_1 increased whereas that of TuTuA_2 decreased (**Figure 5c**). Consistent with our circuit model in females, these results indicate that a decrease in TuTuA_2 activity could relieve LC10a neurons from ongoing inhibition when males become sexually aroused, gating the propagation of visual signals that underlie pursuit behavior. Indeed, activation of TuTuA_2 in males courting a real female increased the average distance between the courting pairs and hindered the ability of males to maintain the female in the center of their field of view (**Supplementary Figure 7g**). These results support the notion that the same TuTuA-mediated switch is used by males and females to gate visual processing during social interactions.

## Concluding Remarks

Animals gate visual information in a context-dependent manner. Using the connectome as a guide, our work provides a detailed circuit level understanding of how this can be accomplished. We found three distinct mechanisms, under coordinated control by a single cell type that conveys internal state, that selectively amplify visual information critical for social interactions (**Figure 6**). This cell type, aIPg, appears to be largely dedicated to this task with its top six synaptic targets contributing to the gating of the visuomotor circuits we described. These circuits are engaged by aIPg and have the potential to regulate distinct motor programs (**Supplementary Figure 9**). The presence of these multiple circuit mechanisms endows the system with more degrees of freedom and flexibility in regulation of attention toward different visual features. It is difficult to imagine how we could have efficiently discovered these circuit mechanisms without the connectome. aIPg is the primary sexually dimorphic neuron in the circuits we described, while other circuit components— the LC neurons, lobula interneurons, IB112 and TuTuA neurons— are present in both males and females with indistinguishable morphologies. Our results illustrate how a single node that differs across sexes could regulate common sensorimotor circuits, indicating that these circuit mechanisms will play a role in a range of social behaviors.

**Fig. 6.**
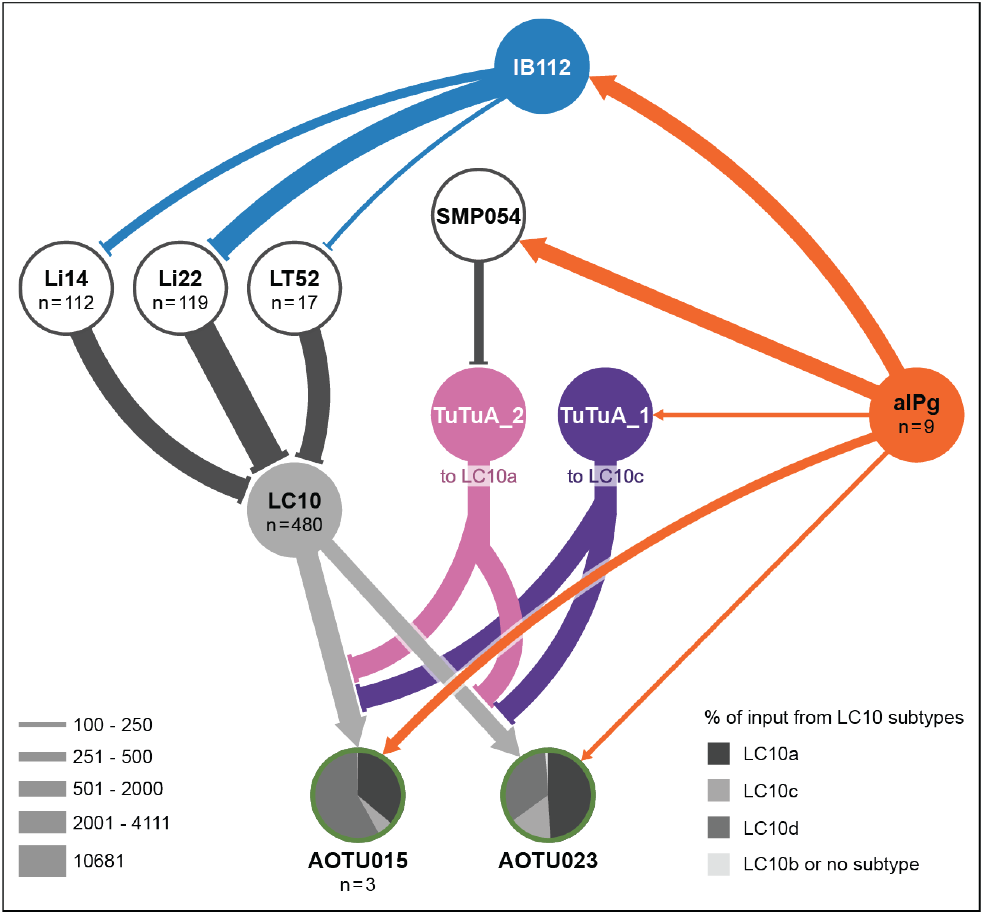
Summary of neural motifs for state-dependent modulation of visual information flow through LC10 neurons. Overview of the circuit components for each mechanism detailed in Figure 1a. Activation of aIPg: (1) provides additional excitatory input to downstream targets of LC10 neurons, represented here by AOTU015 and AOTU023; (2) leads to disinhibition of inputs to the dendrites of LC10 neurons through the action of IB112 on local inhibitory neurons in the optic lobe; and (3) governs whether LC10a or LC10c is able to signal to their downstream targets by a novel toggle switch operated by the TuTuA_1 and TuTUA_2 neurons which provide axo-axonal inhibition to LC10c and LC10a, respectively. See text for details. Line widths represent synaptic connections and are scaled according to the key. For cell types with more than one cell per brain hemisphere, the number of cells are indicated in the circle. See **Supplementary Figure 9** for additional details.

## Acknowledgements

We thank L. Abbott and the Janelia Community for their helpful suggestions during the course of this work and their comments on the manuscript. We also thank the Fly Light team for generating the images of GAL4 expression patterns as well as Karen Hibbard and Heather Dionne for assisting with generation of genetic reagents for female behavioral experiments. This work was supported by Howard Hughes Medical Institute and Simons Foundation Collaboration on the Global Brain, and by an NIH NINDS grant (5R35NS111611) to VR.

## Author Contributions

CES and GMR conceived of and designed the study. GMR, MD, and AN performed connectome analyses. CES performed strain construction as well as female behavioral experiments and analysis. GMR generated cell type specific genetic driver lines with assistance from CES and the Janelia Fly Light Project Team. The construction of the analysis pipeline for this data was designed and developed by KMB, AAR, ALT, and CES. NK performed functional imaging experiments detailed in Fig. 2a and Supplementary Fig. 2a – d. CES and AO performed computational analysis of visual parameters during female aggression and male courtship in Fig. 2b and Supplementary Fig. 2e – f. Electrophysiology recordings and analysis was performed by MS. DB performed in vitro functional imaging experiments shown in Supplementary Fig. 6a – d. THS and VR designed, performed, and analyzed the functional imaging and male courtship experiments in Fig. 5C and male courtship experiments in Supplementary Fig. 7g. MD, CES, THS, and AO prepared graphics for figures. The original draft of the manuscript was written by CES and GMR with input from all authors.

## Methods and Materials

### Fly strains

All experiments used mated female flies unless otherwise stated. Flies were reared on standard cornmeal molasses food at 25°C and 50% humidity. For optogenetic activation experiments, flies were reared in the dark on standard food supplemented with retinal (Sigma-Aldrich, St. Louis, MO) unless otherwise specified, 0.2 mM all trans-retinal prior to eclosion and 0.4 mM all trans-retinal post eclosion. Hemidriver lines were created using gateway cloning as previously described (61). Stable split GAL4 lines used in this study were constructed as described in (61) and hemidrivers used are detailed in the reagents table above. Original confocal image data of GAL4 lines are available at https://www.janelia.org/split-GAL4.

### Thermogenetic and optogenetic activation behavioral experiments

Groups of 5–8 group-housed mated female flies (7–10 days post-eclosion) were video recorded in 60% relative humidity in a 53.3 mm x 3.5 mm circular arena as described in (62). All non-thermogenetic (TrpA) experiments were performed at 24°C, while thermogenetic experiments were performed at 22°C for non-permissive controls and 31°C for permissive tests. All tests were conducted under visible light conditions at ZT0 to ZT4 unless otherwise stated. Flies were loaded into the arena using an aspirator. For activation of neurons expressing CsChrimson, the arena was illuminated as specified in the figure legends using constant uniform illumination with 660 nm LEDs. For inactivation of neurons expressing GtACR, we used constant uniform illumination with 525 nm LEDs. All trials were performed under white-light illumination from above. Videos were recorded from above using a camera (USB 3.1 Blackfly S, Monochrome Camera; Point Gray, Richmond, Canada) with an 800 nm long pass filter (B and W filter; Schneider Optics, Hauppauge, NY) at 170 frames per second and 1024 × 1024 pixel resolution.

Male courtship experiments were performed as detailed in (41). Briefly, all assays were performed with virgin male and virgin female flies 3-6 days post-eclosion. Flies were isolated 2-8 hrs post-eclosion and reared with flies of the same sex at low density. Experiments were performed in custom-milled Delrin chambers (d = 20 mm, h = 2.5 mm or 3.5 mm, protolabs) with sloped edges to decrease the chances of flies walking on edges. A thin layer of transparent acrylic board was used as a lid for the chamber. We added flies to the chamber by aspiration without anesthetization. All assays were performed under bright white light conditions, and red LEDs were used to illuminate the behavioral arena and stimulate ion flux through CsChrimson. Flies expressing CsChrimson were dark-reared on sugar-yeast food (100 g Brewer’s yeast, 50 g sucrose, 15 g agar, 3 ml propionic acid, 3 g p-hydroxybenzoic acid methyl ester per 1 Liter H2O) to avoid low lev-els of retinal metabolized from vitamin A in more nutritious food. Progeny from these crosses were also raised on sugar-yeast food before flies were transferred to food containing 400 *µ*M all-trans-retinal (Sigma-Aldrich R2500) at least 48 hrs before the experiments.

**Table 1.** Aggression and Touch JAABA classifiers.

| Classifier | True Positive | True Negative | False Positive | False Negative |
| --- | --- | --- | --- | --- |
| Aggression | 89.5% (2072) | 98.4% (2515) | 10.5% (244) | 1.6% (40) |
| Touch | 97.4% (1939) | 86.7% (1818) | 2.6% (52) | 13.3% (278) |

### Behavioral classification and analysis

For each trial, flies were acclimatized to the arena for 30 s prior to the delivery of six sets of constant stimuli each lasting 30 s with 30 s in between each stimulus. For all experiments, only the lowest stimulus intensity in which an effect was found is depicted and was analyzed. Unless otherwise stated, the pre-stimulus average was calculated from the three periods prior to the stimulus periods used for analysis. In Figures 2c and Supplementary Figure 7d, the prior stimuli appeared to alter behavior during successive stimulus off periods; therefore, only the first 15 s of the first pre-stimulus period was used for comparison. Flies were tracked using Caltech FlyTracker followed by automated classification of behavior with JAABA classifiers (see (63)). Novel classifiers for touch and aggression were created based on prior definitions (47, 64). We validated the performance of this classifier against manually labeled ground truth data using videos that were not part of the training dataset (see Supplemental table 1 for framewise performance). For figures displaying behavioral time courses, the mean of 2.83 s (60-frame) bins is shown.

For dyadic courtship assays, courtship start frame was manually identified based on first instance inter-fly distance of < 3 mm and fixation angle of < |20°| lasting > 1 minute. Courtship end was defined as first copulation frame or end of video acquisition (30 min).

For calculation of visual features experienced during aggressive and courtship interactions, the angular position, velocity, height, and width were calculated on a frame-by-frame basis using a custom MATLAB (MathWorks) script wherein, for each frame, the coordinates and orientations of subject fly and nearest conspecific, or target fly, were translated and rotated such that the subject was situated at the origin facing zero degrees. In this new basis, the target fly’s angular position (*θ*) and velocity (*φ*) with respect to the subject fly’s visual field were calculated as *θ* `= tan`^−1^`(y/x)` and *φ* = *d**θ*/*dt*, respectively. To approximate angular size and expansion of the target fly in the subject’s visual field, an ellipse was fit to the major and minor axes of the target fly in this new basis. The target fly’s angular width was approximated as the length of the cross-section of this ellipse that lies perpendicular to the Euclidean line between the anterior-most point of the subject fly and target fly centroid. Thus, for each frame, the equation for the target fly’s angular width, *w*, is: *w* = 2(tan^−1^(*R*/d)) where *R* is half the real cross-sectional length of the ellipse (in mm) and *d* is the Euclidean distance (in mm) between the anterior-most point of the subject fly and target fly centroid. The target fly’s angular height was approximated in a similar manner, however the target fly’s real height was fixed at 1mm (which a reasonable estimation of a female fly’s height), so the equation for angular height, *h*, was simply: *h* = 2(tan^−1^(*0*.*5 mm*/*d*)) where *d* is the Euclidean distance (in mm) between the anterior-most point of the subject fly and target fly centroid.

### Fly preparation for calcium imaging

For Figure 2 and Supplementary Figure 2a – c, experiments were performed similar to (23) with a similar preparation as described in (65). Notably, the fly’s head was positioned and glued to the fly holder such that the eye’s equator faced the middle of the visual projection screen. The proboscis remained intact but was glued in position, and a dissection needle was used to remove the cuticle and sever muscle 16.

For Figure 5, experiments were performed similarly to (41). Briefly, flies were anesthetized on *CO*_2_ and tethered to a custom-milled plate. Flies were held in place by a string across the neck and fixed to the holder by both eyes and the back of the thorax using UV-curable glue. The proboscis was also glued to the mouthparts to minimize brain motion. Flies were left to recover in a warm, humidified chamber (25°C, 50-70% humidity) in the dark for 1-4 hrs. The cuticle was subsequently dissected from the top of the head and flies were transferred to an air-supported foam ball.

### Two-photon calcium imaging

In Figure 2 and Supplementary Figure 2a – c, calcium imaging experiments were performed with LC10a male flies or LC9 female flies 5-10 days post-eclosion, maintained under standard conditions (21.8°C, 55% humidity, 16 hr light/8 hr dark, standard corn-meal and molasses food). The key resources table lists fly genotypes used in calcium imaging experiments. The imaging setup is identical to the previously described two-photon microscope (Thorlabs) setup (23). Briefly, we used a Ti:Sapphire femtosecond laser (Spectra-Physics Mai Tai eHP DS) tuned to 920 nm and delivering <20 mW power at the sample. Fluorescence signals were collected using a 16x water-immersion objective (Nikon CFI75, NA 0.8) with a bandpass filter (Semrock 503/40 nm) in front of the photomultiplier tube (Hamamatsu GaAsP H10770PB-40 SEL). Oxygenated saline was circulated throughout. Imaging volumes were acquired at 5.6 Hz or higher. Visual stimuli were delivered to the fly’s right eye and all imaging was from the right side of the brain. The stimuli were presented on a screen that subtended roughly ≈90° by ≈90° of the fly’s field of view with a green (532 nm) projector setup as previously described (23).

In Figure 5, calcium imaging experiments were performed with TuTuA_1 and TuTuA_2 male flies 3-7 days post-eclosion, maintained under standard conditions (25°C, 65% humidity, 12 hr light/12 hr dark, standard Würzburg food).

The imaging preparation for tethered courtship was identical to that previously described in (41). Briefly, male flies rested and walked on a small 6.35 mm diameter ball, which was shaped from foam and manually painted with uneven black spots using a Sharpie. The foam ball was held by a custom-milled aluminum base and floated by air supplied at ≈0.8 L/min such that the ball could move smoothly. The ball was illuminated by infrared LED flood lights, and imaged with a Point Grey FLIR Firefly camera by way of a mirror. The ball was surrounded by a 270° conical screen with a large diameter of ≈220 mm, a small diameter of ≈40 mm, and a height of ≈60 mm. As males walked on the foam ball, all three rotational axes of the ball were read out by the FicTrac2.0 software (66) at 60 Hz in real-time. The visual stimulus was projected around the male from a DLP 3010 Light Control Evaluation Module (Texas Instruments), via a first-surface mirror below the fly. The red and green LEDs in the projector were turned off, leaving only the blue LEDs to minimize interference with GCaMP emissions.

Visual stimuli were generated in the MATLAB-based ViRMEn (67) software and projected onto the screen using custom perspective transformation functions. The net visual refresh rate of the visual stimulus ranged from 47.6 Hz to 58.9 Hz. Each trial was initiated by the presentation of a stationary visual target for 60 s to examine the animal’s baseline locomotion, after which the visual target began to oscillate. The visual target oscillated in a 107° arc around the animal with a constant angular velocity of ≈75°/s, but the angular size of the dot was continuously altered to mimic the dynamics of a natural female during courtship. The angular size was altered by changing the distance between the male and the target in the ViRMEn world. The distance between the male and the target was taken from the inter-fly-distance (IFD) in a courting pair over the course of two minutes of courtship, and at each frame, the angular position of the target was scaled by this IFD to give rise to a more dynamic female path. Angular sizes ranged between ≈8-50°, with the average size being 22.5°. Each stimulus frame was thus unique for 2 min of time, and subsequently repeated until the end of the trial when it intersected its original position. Each trial lasted 10 min.

Male imaging experiments were performed with an Ultima Investigator or Ultima Investigator Plus two-photon laser scanning microscope (Bruker Nanosystems) with a Chameleon Ultra II Ti:Sapphire laser. All samples were excited at a wavelength of 920 nm, and emitted fluorescence was detected with a GaAsP photodiode detector (Hamamatsu). All images were acquired with a 40X Olympus water-immersion objective with 0.8 NA. All images were collected using PrairieView Software (Version 5.5 or 5.7) at 512 pixel x 512 pixel resolution

Courtship and running was classified based on the fidelity and vigor of a male’s pursuit of the visual target, as described in (41).

In Supplementary Figure 6, ex vivo calcium imaging experiments were performed similar to (68). Briefly, flies were reared at 25°C on cornmeal medium supplemented with retinal (0.2 mM) that was shielded from light. All experiments were performed on female flies, 3–5 days post-eclosion. Brains were dissected in a saline bath (103 mM NaCl, 3 mM KCl, 2 mM CaCl2, 4 mM MgCl2, 26 mM NaHCO3, 1 mM NaH2PO4, 8 mM trehalose, 10 mM glucose, 5 mM TES, bubbled with 95% O2/5% CO2). After dissection, the brain was positioned anterior side up on a coverslip in a Sylgard dish submerged in 2 ml saline at 20°C. The sample was imaged with a resonant scanning 2-photon microscope with near-infrared excitation (920 nm, Spectra-Physics, INSIGHT DS DUAL) and a 25X objective (Nikon MRD77225 25XW). The microscope was controlled by using ScanImage 2017 (Vidrio Technologies). Volumes were acquired with 230 *µ*m × 230 *µ*m field of view at 512 × 512 pixel resolution at 2 *µ*m steps over 42 slices, at approximately 1 Hz. The excitation power for Ca^2+^ imaging measurement was 15 mW. On the emission side, the primary dichroic was Di02-R635 (Semrock), the detection arm dichroic was 565DCXR (Chroma), and the emission filters were FF03-525/50 and FF01-625/90 (Semrock). During photostimulation, the light-gated ion channel CsChrimson was activated with a 660 nm LED (M660L3 Thorlabs) coupled to a digital micromirror device (Texas Instruments DLPC300 Light Crafter) and combined with the imaging path with a FF757-DiO1 dichroic (Semrock). Photostimulation occurred at 10Hz over two periods with a duration of 14 s at 0.037 mW/mm^2^ intensity interspersed by a 2 s pause. After responses to the photostimulation, laser power was increased to take two color highresolution images containing fluorescence from both red and green channels. Using custom python scripts, ROIs corresponding to cell compartments were identified in the high resolution images. These ROIs were then applied to the time series images to measure intensity changes in response to the photostimulation. Fluorescence in a background ROI, that contained no endogenous fluorescence, was subtracted from the cell compartment ROIs. In the ΔF/F calculations, baseline fluorescence is the mean fluorescence over a 10 s time period before stimulation started. The ΔF is the fluorescence minus the baseline. Then the ΔF is divided by baseline to normalize the signal (ΔF/F).

### Electrophysiology

Whole-cell patch-clamp recordings were obtained from freshly isolated brains of 3–5 day old flies. The brain was continuously perfused with oxygenated (95% O2/5% CO2) extracellular saline containing (in mM): 103 NaCl, 3 KCl, 1.5 CaCl2·2H2O, 4 MgCl2·6H2O, 1 NaH2PO4·H2O, 26 NaHCO3, 5 TES, 10 Glucose, and 10 Trehalose·2H2O. Osmolarity was 275 mOsm, and pH was 7.3. Recording electrodes were pulled from thick-walled glass pipette (1.5 mm/0.86 mm) using P-97 puller (Sutter Instruments) and fire polished using MF 830 (Narishige) to achieve resistances of 10–12 MΩ. Intracellular saline contained (in mM): 137 KAsp, 10 HEPES, 1.1 EGTA, 0.1 CaCl2·2H2O, 4 MgATP, 0.5 NaGTP. Osmolarity was 260-265 mOsm, and pH was adjusted to 7.3 with KOH. Biocytin was added to intracellular solution at 0.5% for post hoc morphological confirmation.

The brain was visualized by an IR-sensitive CCD cam-era (ThorLabs 1501M) with an 850 nm LED (ThorLabs M850F2). GFP-labeled cell body was visualized with 460 nm LED (Sutter Instruments). Images were acquired using Micro-Manager with automatic contrast adjustment. Recordings were obtained from cell bodies under a 60× water-immersion objective (Olympus).

Current-clamp recordings were sampled at 20 KHz, low-pass filtered at 10 KHz using Digidata 1550B, Multiclamp 700B, and Clampex 11.2 software (Molecular Devices). Recordings were made at membrane potential of -50 mV to -65 mV, with small (5-30 pA) hyperpolarizing current injections as needed, and not corrected for liquid junction potentials.

CsChrimson was activated by 630 nm LED at 0.4 mW/cm^2^. Stimulation duration was set at minimal value which is sufficient to induce reliable responses from target neurons. After the electrophysiology recording, the whole brain was fixed in 4% paraformaldehyde in 0.1 M PBS until further staining. After rinsing in PBS, the brain was incubated in Streptavidin Alexa Fluor 647 (1:200) in PBS-T overnight at room temperature. The preparations were then rinsed, dehydrated and mounted with DPX. The confocal images were captured with LSM 980 microscope (Zeiss), with 639 nm excitation wavelength.

The electrophysiological recordings were analyzed using pClamp (Clampfit 11.3). The instantaneous action potential frequency was calculated for about one minute in each cell. The action potential amplitude was averaged from 20-30 individual events in each cell, and measured as the difference between the threshold and peak.

### Immunohistochemistry and imaging

All experiments were performed as described previously (47, 69–73). Additional details of the imaging pipeline used are available at https://data.janelia.org/pipeline.

### Connectomics analysis

Our analyses are based on the hemi-brain (24) dataset (v1.2.1) as queried using the neuPrint interface (neuprint.janelia.org) unless otherwise noted. The unique identifier (bodyID number in the hemibrain v1.2.1 database) for neurons are shown in figures, and a complete list of synaptic connections used to construct our circuit diagrams can be found in neuPrint. Because the hemibrain did not include the entire lobula we also performed analyses in the recently completed and fully annotated male optic lobe connectome (54). These analyses were then confirmed with those observed in the Flywire (55) analysis of the FAFB dataset (74) queried using Codex (https://codex.flywire.ai). Similarly, connections to DNs described in Extended Figure 9 were evaluated using Flywire.

### Statistics

No statistical methods were used to pre-determine sample size. Sample size was based on previous literature in the field and experimenters were not blinded in most conditions as all data acquisition and analysis were automated. Biological replicates completed at separate times using different parental crosses were performed for each of the behavioral experiments. Behavioral data are representative of at least two independent biological repeats. For figures in which the behavioral data over the course of a trial is shown, a yellow or red bar indicates the stimulus period, the mean is represented as a solid line, and shaded error bars represent variation between experiments.

For each experiment, the experimental and control flies were collected, treated and tested at the same time. A Wilcoxon Matched-pairs Signed Rank test (two-tailed) was used for statistical analysis of optogenetic experiments when examining effects within the same group. For analysis among two groups, a Mann-Whitney test (two-tailed) was used, while a Kruskal-Wallis test with Dunn’s multiple comparisons posthoc analysis was used to compare across multiple groups. All statistical analysis was performed using Prism Software (GraphPad, version 10). p values are indicated as follows: ****p<0.0001; ***p<0.001; **p<0.01; and *p<0.05. See Supplementary file 1 for exact p-values for each figure.

For bar plots, all data points are shown to indicate the range and top edge of bar represents the mean. Boxplots show median and interquartile range (IQR). Lower and upper whiskers represent 1.5 × IQR of the lower and upper quartiles, respectively; boxes indicate lower quartile, median, and upper quartile, from bottom to top. When all points are shown, whiskers represent range and boxes indicate lower quartile, median, and upper quartile, from bottom to top. In violin plots, lower and upper quartiles are indicated with dotted light grey lines, while median is indicated with a solid light grey line. Shaded error bars on graphs are presented as mean ± s.e.m.

---

**Fig. S1.**
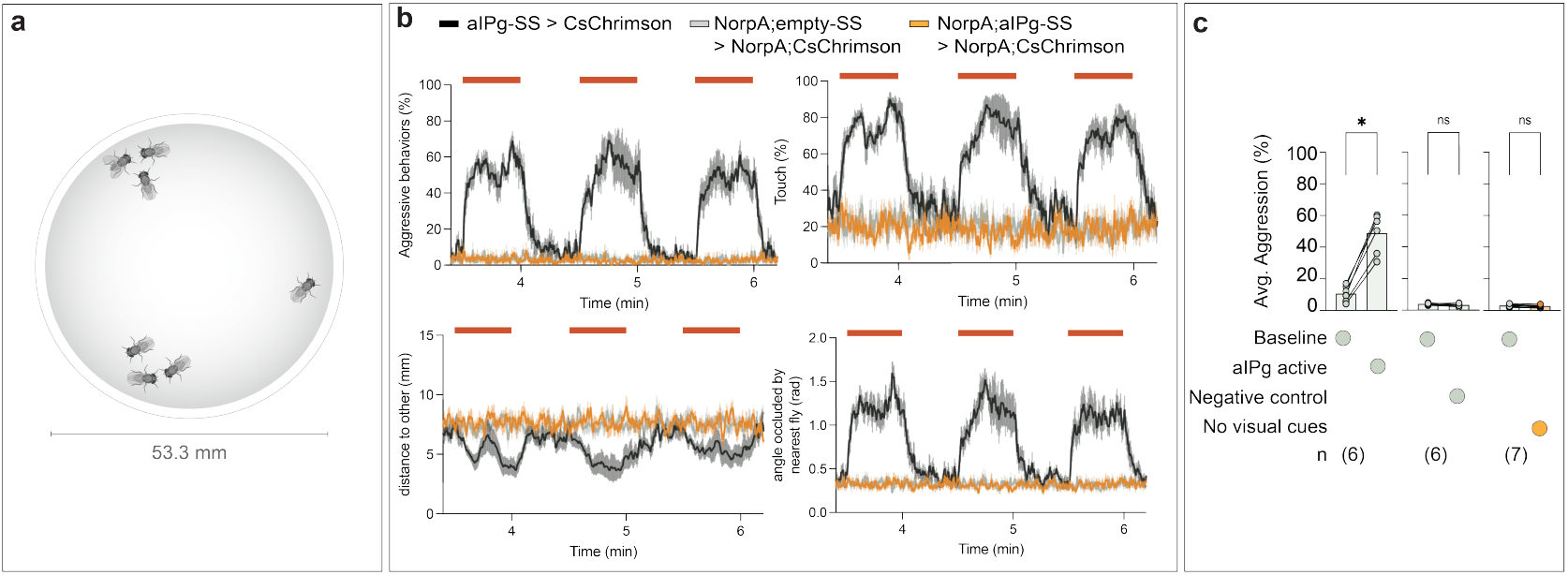
Pathways carrying visual information are important for female aggressive behaviors and related behavioral features. (a) Diagram of the arena used for female behavioral experiments. We performed behavioral experiments in a standardized 53.3 mm arena in which freely moving fly behavior was quantified using a 170 frames per second camera and computer vision-based classification methods (62, 64). (b) Percentage of flies engaging in aggressive behaviors, touching, and changes in parameters related to distance to another fly and the maximum angle of the field of view occluded by the closest fly (angle occluded by nearest fly) are plotted over the course of a 3.3 min trial during which a 3x 30 s 3 mW/cm^2^ continuous red-light stimulus (red bars) were delivered. Prior data are not shown as no significant changes were found in the no visual cues group during this low stimulus period (1 mW/cm^2^) as well. The mean is represented as a solid line and shaded bars represent standard error between experiments. The timeseries shows the percentage of flies performing aggression displayed as the mean of 2.83 s (60-frame) bins. (c) Average time spent performing aggressive behaviors before and during stimulus periods. All data points are shown to indicating the range and top edge of bar represents the mean. Each dot represents one experiment containing approximately seven flies. Data supporting the plots shown in panels b – c were as follows: aIPg-SS > CsChrimson, n = 6 experiments; norpA, EmptySS > norpA, CsChrimson, n = 6 experiments; norpA, aIPg-SS > norpA, CsChrimson, n = 7 experiments. Data are representative of two independent replicates, which included separate parental crosses and were collected on different days. A non-parametric Wilcoxon Matched-pairs Signed Rank test was used for statistical analysis. Asterisk indicates significance from 0: *p<0.05.

**Fig. S2.**
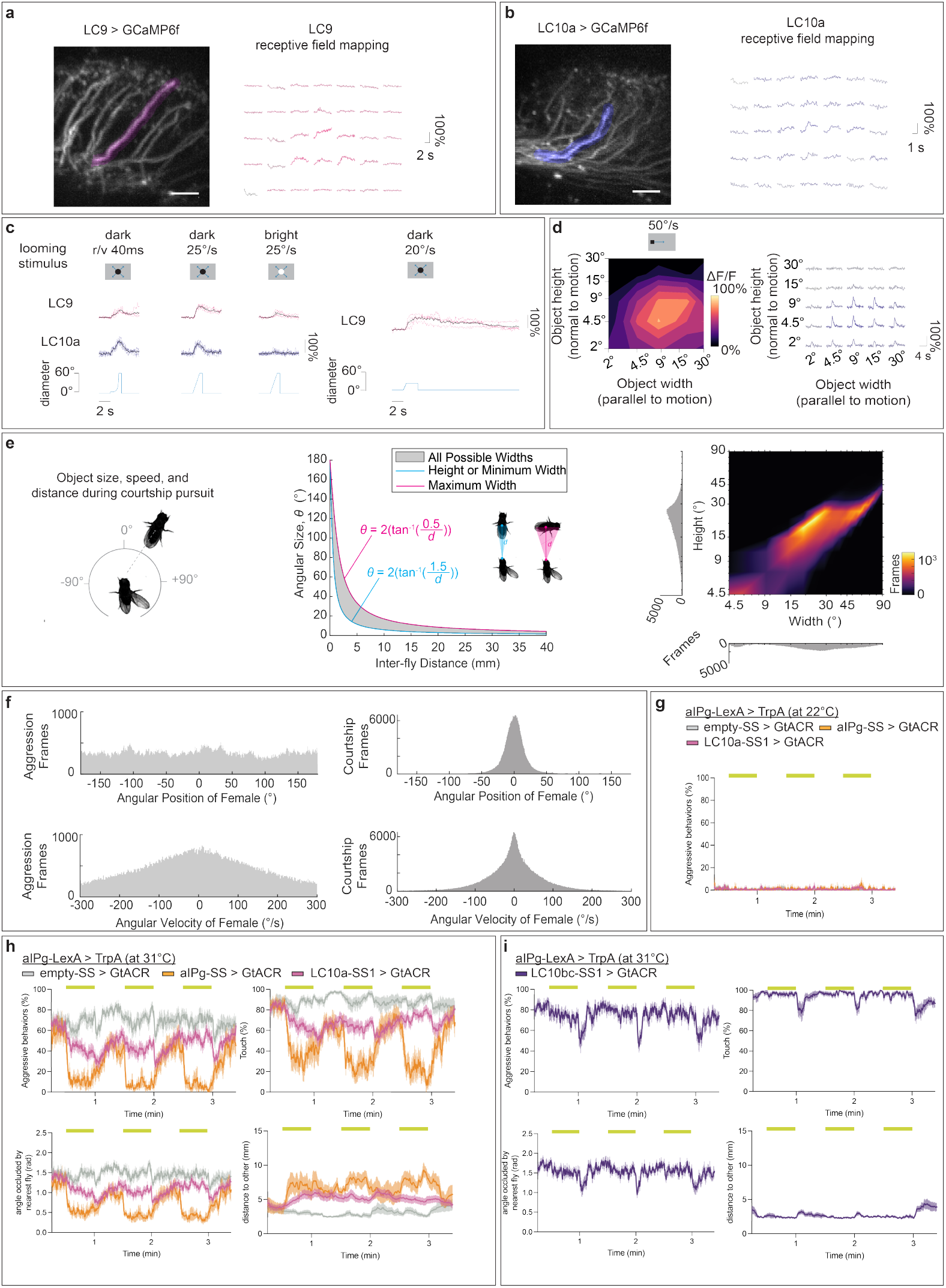
LC visual feature detection and involvement in female aggression. Panels a – d show the receptive fields for LC9 and LC10a. Panels e – f show the visual experience during male courtship and female aggression. Panels g – i demonstrate the effects of inhibiting LC10a or LC10bc during aIPg activation. (a – b) Single receptive-field mapping for individual LC axons in representative flies. LC9 and LC10a axonal regions of interest are colored in magenta and blue, respectively and overlaid on averaged calcium image (left; Scale bar: 10 *µ*m). Individual calcium responses, arranged as in **Figure 2a**, are shown on right. (c) Single-cell (color) and population average (black) calcium traces for neurons responding to looming stimuli centered on the receptive field, same as performed in (23). LC9: n = 4 flies, n = 4 neurons. LC10a: n = 5 flies, n = 7 neurons (25°/s constant edge speed looming was only recorded for 2 neurons from 1 fly). (D) Size tuning, as measured and plotted in **Figure 2a**, for objects of varying sizes moving at a slower speed of 50°/s. (e – f) (e) Left: histograms show conspecific angular position in the visual field as experienced by the male during courtship pursuit. Visual parameters were calculated from single choice courtship assays (n = 13). During male courtship pursuit, the mean conspecific size as subtended on the retina was 15.96 +/- 4.4° (mean +/- standard deviation) in height and 28.699 +/- 12.7° in width. Right: all possible angular heights and widths for a female with a minor axis of 1 mm and major axis of 3 mm are plotted on the left, and the measured angular sizes and heights across frames in the courtship assays are shown in heat map and histogram representations on far right. (f) Histograms of angular position and velocity during during female aggression (left) and naturalistic male courtship pursuit (right). (g – i) Percentage of flies engaging in behaviors (aggression, touch) or behavioral features (distance to other, angle occluded by nearest fly) over the course of a trial during which 3x 30 s continuous green light (yellow bars) were delivered. To control for additional cell types in the LexA line used for aIPg (75), we simultaneously inhibited aIPg during thermogenetic activation through using an aIPg-specific split-GAL4 line and the green light gated anion channel, GtACR. The dramatic reduction in female aggressive behavior during optogenetic inhibition confirmed that aIPg was primarily responsible for the aggression observed when stimulating the LexA line. Data supporting the plots shown in panels g – i were as follows: g: aIPg-LexA > TrpA emptySS > GtACR, n = 10 experiments; aIPg-LexA > TrpA aIPg-SS > GtACR, n = 8 experiments; aIPg-LexA > TrpA LC10a-SS > GtACR, n = 13 experiments. h: aIPg-LexA > TrpA emptySS > GtACR, n = 15 experiments; aIPg-LexA > TrpA aIPg-SS > GtACR, n = 8 experiments; aIPg-LexA > TrpA LC10a-SS > GtACR, n = 18 experiments. i: aIPg-LexA > TrpA LC10bc-SS > GtACR, n = 11 experiments. Experiments were performed at the permissive temperature (31°C, h – i) for aIPg > TrpA stimulation, and non-permissive temperature controls (22°C) are shown in g. The mean is represented as a solid line and shaded bars represent standard error between experiments. The timeseries shows the percentage of flies performing aggression displayed as the mean of 2.83 s (60-frame) bins.

**Fig. S3.**
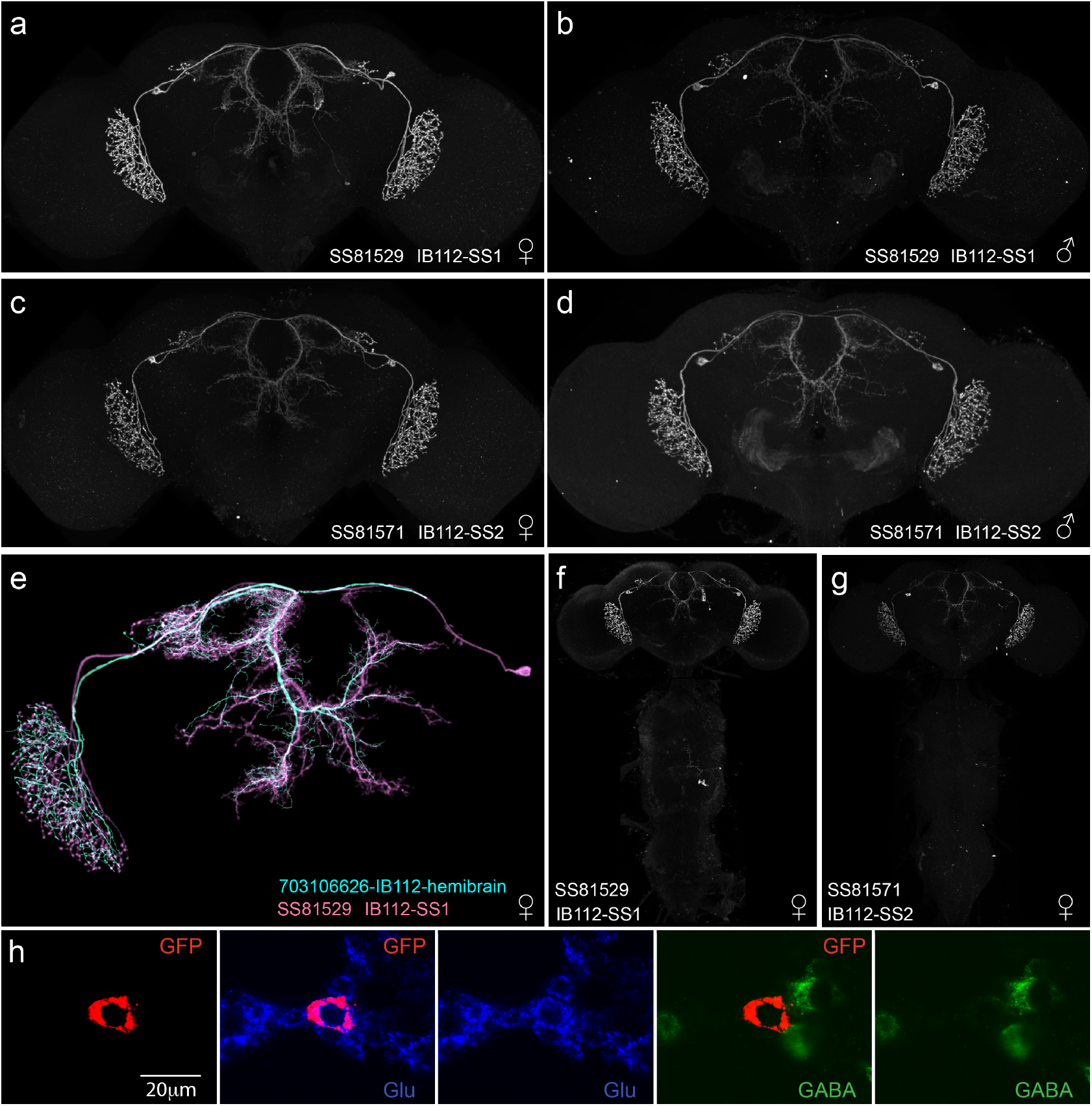
Anatomy of GAL4 driver lines for IB112. (a, b) Expression patterns in a female and male brain, respectively, of GAL4 line SS81529 (IB112-SS1). (c, d) Expression patterns in a female and male brain, respectively, of GAL4 line SS81571 (IB112-SS2). (e) IB112 body ID 703106626 skeleton from hemibrain v1.2.1 shown together with a neuron from SS81529 obtained by stochastic labeling (70) and then segmented using VVD (see Key Resources table). (f, g) Images of the expression patterns in the brain and VNC of GAL4 driver lines SS81529 and SS81571, as indicated. (h) Images of fluorescent in situ hybridization assays to determine the neurotransmitter used by IB112. Probes used in each panel are indicated. GFP shows the IB112 cell body and Glu and GABA represent probes for vGlut and GAD, respectively (see Methods for details). Scale bar is shown in the left panel.

**Fig. S4.**
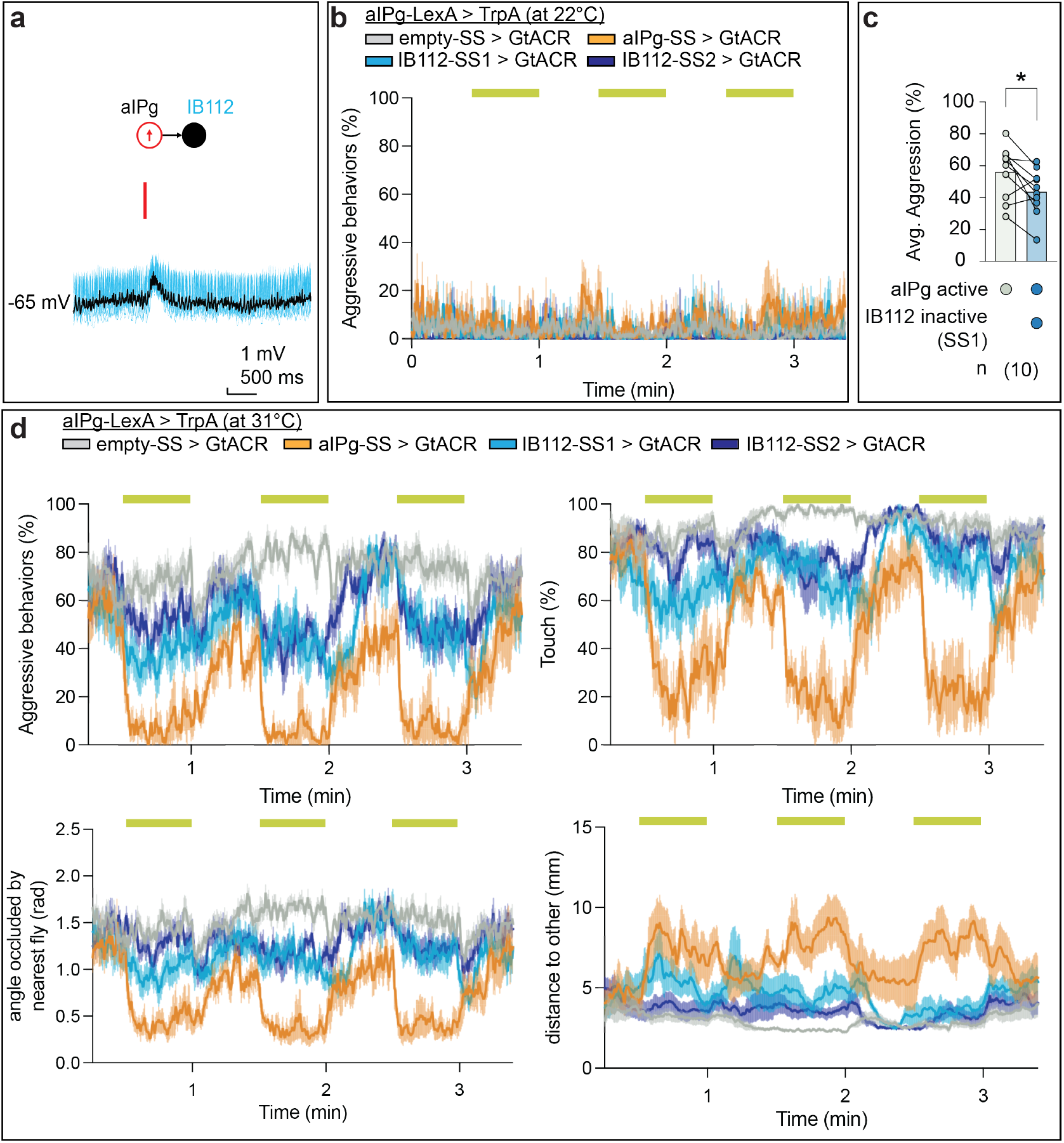
IB112 shapes aIPg-mediated female aggressive behaviors. (a) Excitatory responses recorded by patch clamp electrophysiology in female brain explants from IB112 (n = 6 cells) before, during, and following a 15 ms activation of aIPg. Individual trials in blue (n = 8 trials from one cell), mean shown in black. (b, d) Percentage of flies engaging in aggression, touch, or changes in related parameters, including the maximum angle of the field of view occluded by the closest fly (angle occluded by nearest fly) or distance to another fly. Percentages are plotted over the course of a trial during which 3x 30 s 9 mW/cm^2^ continuous light stimuli (yellow bars) were delivered. The mean is represented as a solid line and shaded bars represent standard error between experiments. The timeseries shows the percentage of flies performing aggression displayed as the mean of 2.83 s (60-frame) bins. (c) Average time spent performing aggressive behaviors before and during stimulus periods. Averages were calculated over all flies in an experiment, and each dot represents one experiment containing approximately seven flies. All data points are shown to indicating the range and top edge of bar represents the mean. Data supporting the plots shown in panels b – d were as follows: b: aIPg-LexA > TrpA emptySS > GtACR, n = 11 experiments; aIPg-TrpA > CsChrimson aIPg-SS > GtACR, n = 5 experiments; aIPg-TrpA > CsChrimson IB112-SS1 > GtACR, n = 4 experiments; aIPg-TrpA > CsChrimson IB112-SS2 > GtACR, n = 4 experiments. c – d: aIPg-LexA > TrpA emptySS > GtACR, n = 13 experiments; aIPg-TrpA > CsChrimson aIPg-SS > GtACR, n = 6 experiments; aIPg-TrpA > CsChrimson IB112-SS1 > GtACR, n = 10 experiments; aIPg-TrpA > CsChrimson IB112-SS2 > GtACR, n = 11 experiments. Experiments were performed at the permissive temperature (31°C, c – d for aIPg > TrpA stimulation, and non-permissive temperature controls (22°C) are shown in b. Data were pooled from four independent replicates, which included separate parental crosses and were collected on different days. A non-parametric Wilcoxon Matched-pairs Signed Rank test was used for statistical analysis. Asterisk indicates significance from 0: *p<0.05.

**Fig. S5.**
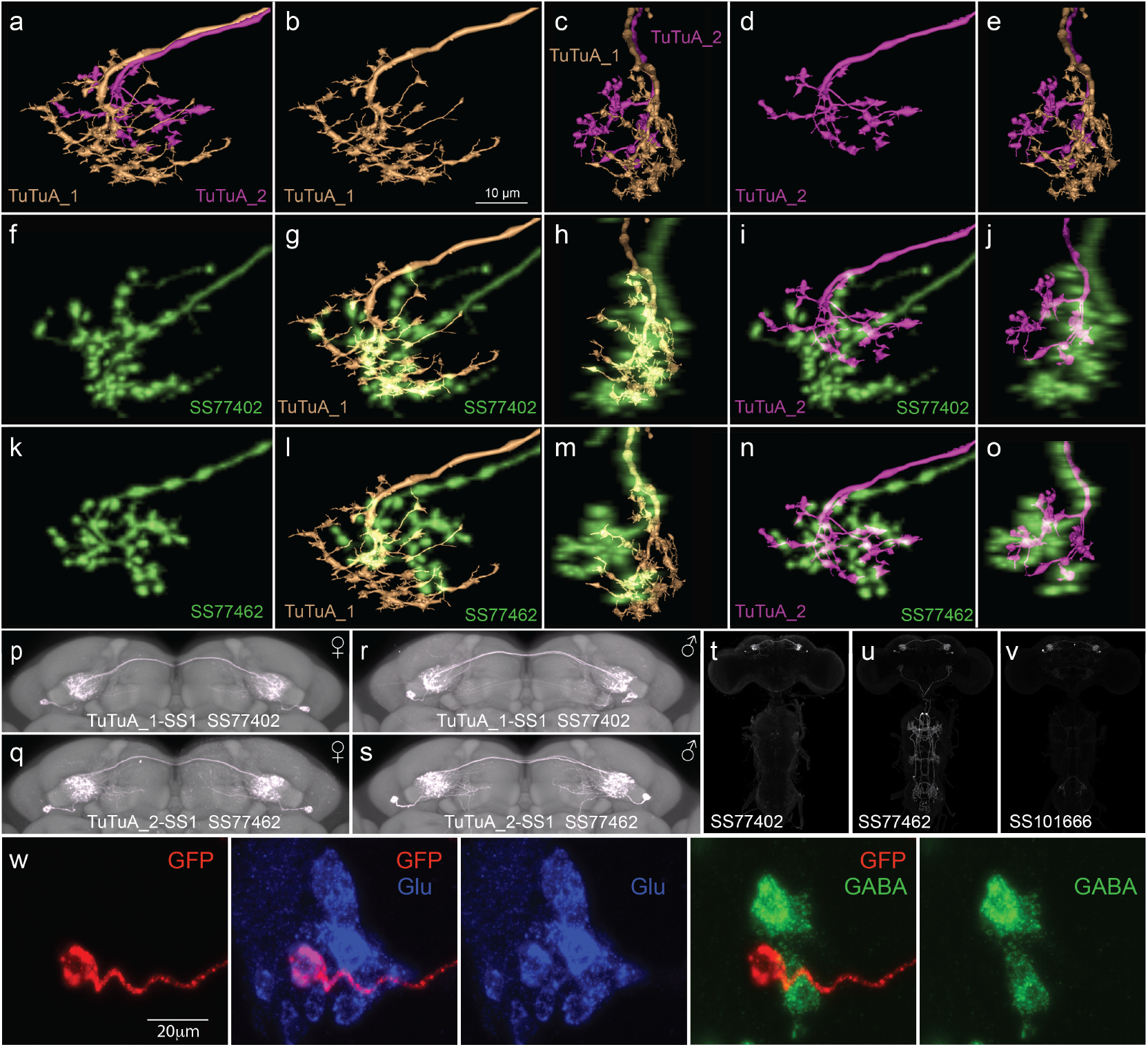
Anatomy of TuTuA subtypes. (a – e) Neuronal skeletons of the termini of the contralateral axons of TuTuA subtypes from the hemibrain v1.2.1 connectome. (a) TuTuA_1 and TuTuA_2 are shown together (body IDs 676836779 and 5813013691, respectively). (b) TuTuA_1 (body ID 676836779) shown alone. (c) Same as panel a, but rotated 90 degrees along the medial-lateral axis. (d) TuTuA_2 (body ID 5813013691) shown alone. (e) Same as panel c, repeated to facilitate comparison. These anatomical differences were used to determine the correspondence between GAL4 driver lines and TuTuA subtypes. (f) Terminus of a contralateral axon of a neuron from GAL4 driver line SS77402 obtained by stochastic labeling(70) and then segmented using VVD (see Key Resources table). (g) The comparison between the GAL4 driver line in f to TuTuA_1 skeleton shown in b. (h) Same as panel g, but rotated 90 degrees along the medial-lateral axis. (i) The comparison between the GAL4 driver line in f to TuTuA_2 skeleton shown in d. (j) Same as panel i but rotated 90 degrees along the medial-lateral axis. (k) Terminus of a contralateral axon of a neuron from GAL4 driver line SS77462 obtained by stochastic labeling(70) and then segmented using VVD. (l) The comparison between the GAL4 driver line in k to TuTuA_1 skeleton shown in b. (m) Same as panel g, but rotated 90 degrees along the medial-lateral axis. (n) The comparison between the GAL4 driver line in k to TuTuA_2 skeleton shown in d. (o) Same as panel i, but rotated 90 degrees along the medial-lateral axis. (p, r) Images of GAL4 driver line SS77402 in females and males, respectively, shown with the standard neuropil reference, JFRC2018U (76) (q, s) Images of GAL4 driver line SS77462 in females and males, respectively. Note the presence of a single TuTuA cell body in each brain hemisphere. (t – v) Images of the expression patterns in the brain and VNC of GAL4 driver lines SS77402, SS77462 and SS10166, as indicated. (w) Images of fluorescent in situ hybridization assays to determine the neurotransmitter used by TuTuA. Probes used in each panel are indicated. GFP shows the TuTuA cell and Glu and GABA represent probes for vGlut and GAD, respectively (see Methods for details). Scale bar is shown in the left panel.

**Fig. S6.**
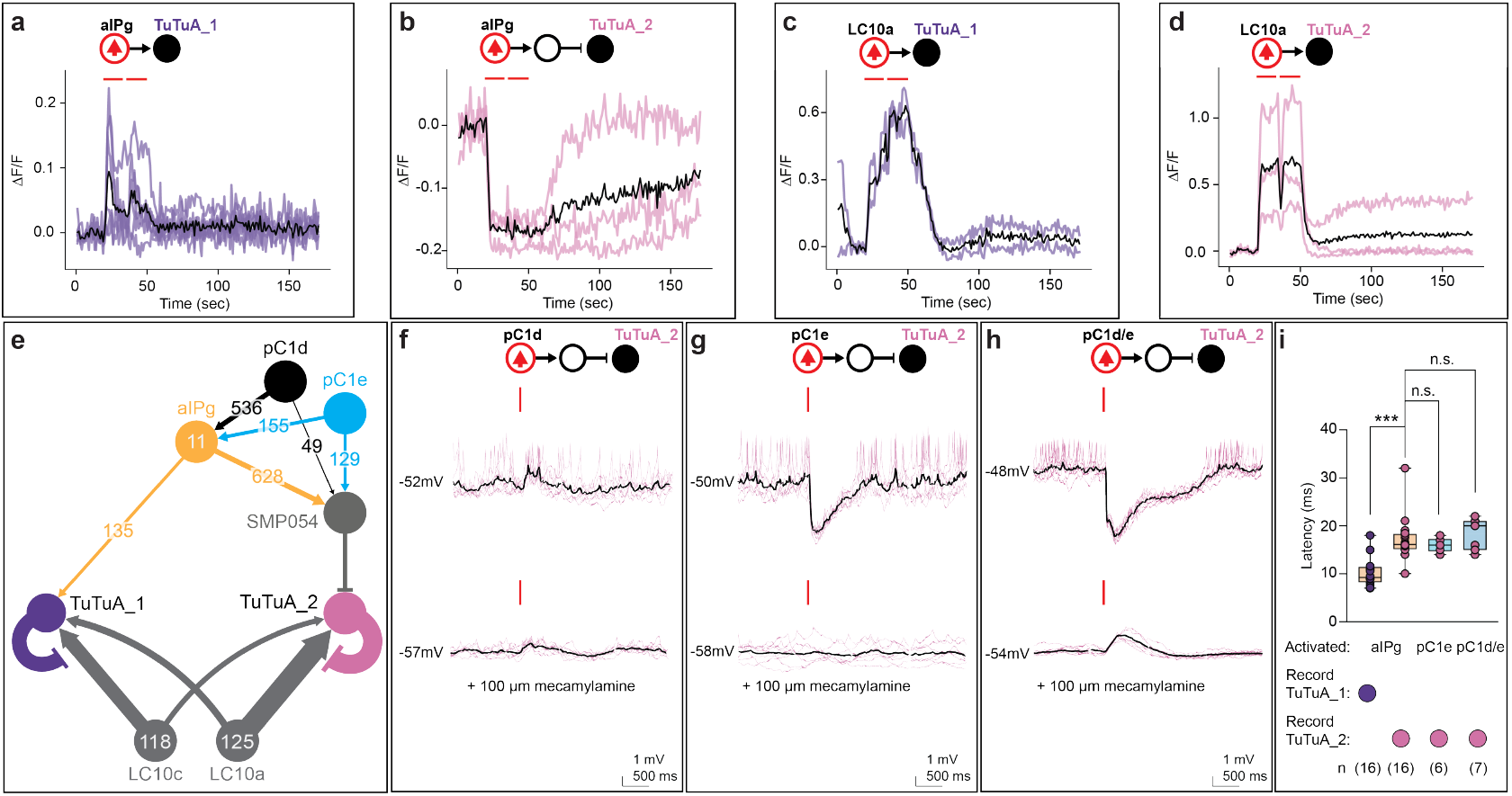
Responses of TuTuA subtypes to the activation of female aggression or LC10a cell types. (a – d) Changes in florescence intensity as measured by GCaMP6f in the cell body of TuTuA_1 (a, c) or TuTuA_2 (b, d) before, during, and following two 14 s stimuli (2 s interval) at 10 Hz. Individual trials for a – d are shown in purple (TuTuA_1) or pink (TuTuA_2), mean is in black. (e) Connectivity diagram from **Figure 4a** with the connections from pC1d and pC1e. Exact synapse numbers are indicated on the arrows, which are also scaled according to synapse counts. Arrows indicate putative excitatory connections (cholinergic) and bar endings indicate putative inhibitory connections (SMP054, GABAergic; TuTuA_1 and TuTuA_2, glutamatergic). (f – h) Electrophysiology recordings with the cell types activated with CsChrimson are circled in red, and those recorded are in black. Individual trials are in pink (n = 8 trials from 1 cell), mean is in black. (f) Small excitation or negligible response in TuTuA_2 (n = 5 cells) to 15 ms pC1d activation, which was abolished by mecamylamine. (g) Large inhibitory response in TuTuA_2 to 15 ms pC1e activation, which was abolished by mecamylamine (n = 6 cells). (h) Large inhibitory response in TuTuA_2 to 15 ms pC1d/e activation, which was abolished by mecamylamine (n = 7 cells). (i) Latency after stimulus onset (ms). Box-and-whisker plots show median and IQR; whiskers show range. A Kruskal-Wallis and Dunn’s post hoc test was used for statistical analysis. Asterisk indicates significance from 0: ***p<0.001.

**Fig. S7.**
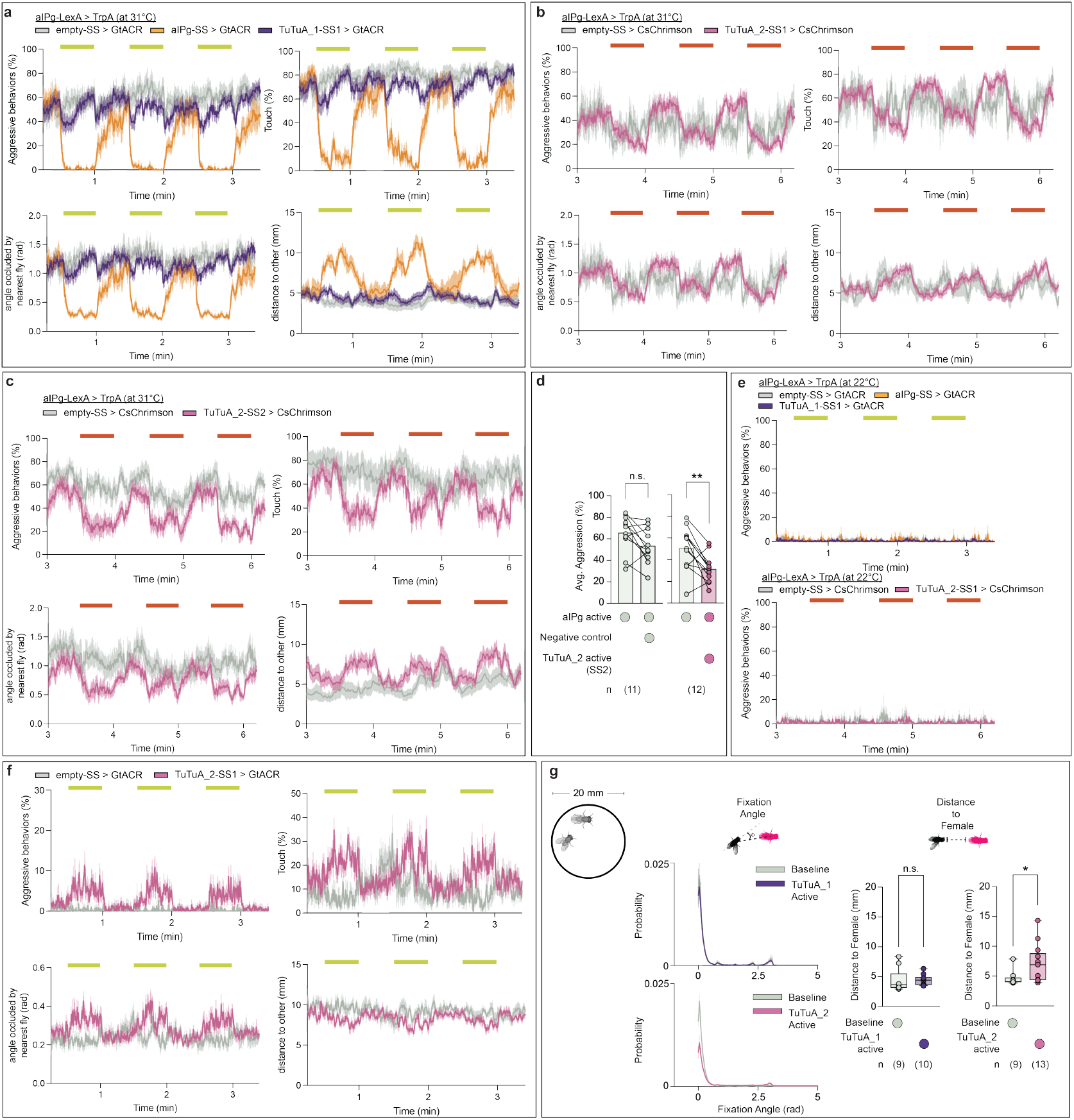
The TuTuA switch shapes female aggression and male courtship behaviors. (a – c, e – f) Percentage of flies engaging in behaviors (aggression, touch) or behavioral features (distance to other, angle occluded by nearest fly) over the course of a trial during which 3x 30 s continuous light stimuli (yellow or red bars) were delivered. Experiments were performed at the permissive temperature (31°C, a – c) for aIPg > TrpA stimulation, and non-permissive temperature controls (22°C) are shown in e. (d) Percentage of flies engaging in aggression over the course of a 3.3 min trial during which 3x 30 s continuous 9 mW/cm^2^ green light (yellow bars) were delivered. Data were pooled from three independent replicates, which included separate parental crosses and were collected on different days. Data supporting the plots shown in panels a – f were as follows: a: aIPg-LexA > TrpA emptySS > GtACR, n = 12 experiments; aIPg-LexA > TrpA aIPg-SS > GtACR, n = 7 experiments; aIPg-LexA > TrpA TuTuA_1-SS > GtACR, n = 20 experiments. b: aIPg-LexA > TrpA emptySS > CsChrimson, n = 6 experiments; aIPg-LexA > TrpA TuTuA_2-SS1 > CsChrimson, n = 18 experiments. c, d: aIPg-LexA > TrpA emptySS > CsChrimson, n = 11 experiments; aIPg-LexA > TrpA TuTuA_2-SS2 > CsChrimson, n = 12 experiments. e (top panel): aIPg-LexA > TrpA emptySS > GtACR, n = 11 experiments; aIPg-LexA > TrpA aIPg-SS > GtACR, n = 4 experiments; aIPg-LexA > TrpA TuTuA_1-SS, n = 11 experiments. e (bottom panel): aIPg-LexA > TrpA emptySS > CsChrimson, n = 4 experiments; aIPg-LexA > TrpA TuTuA_2-SS1 > CsChrimson, n = 9 experiments. f: emptySS > GtACR, n = 6 experiments; TuTuA_2-SS1 > GtACR, n = 11 experiments. The mean for a – c and d - f is represented as a solid line and shaded bars represent standard error between experiments. The timeseries shows the percentage of flies performing aggression displayed as the mean of 2.83 s (60-frame) bins. For figures b – c and the bottom panel of e, data from the low stimulus periods (1 mW/cm^2^) prior are not shown as no significant changes were found. Averages were calculated over all flies in an experiment, with each dot representing one experiment containing approximately seven flies. All data points are shown to indicating the range and top edge of bar represents the mean. (g) Facing angle and average distance to female flies during a male-female pair courtship assay. Diagram of the arena used for courtship experiments shown in inset image on the left. The following genotypes were used: (Left panel) Baseline (TuTuA_1-SS1 > CsChrimson, without retinal), TuTuA_1 active (TuTuA_1-SS1 > CsChrimson, with retinal); (Right panel) Baseline (TuTuA_2-SS1 > CsChrimson, without retinal), TuTuA_2 active (TuTuA_2-SS1 > CsChrimson, with retinal). Box-and-whisker plots show median and IQR; whiskers show range. A non-parametric Wilcoxon Matched-pairs Signed Rank test (d) or Mann-Whitney test (g) was used for statistical analysis and each dot represents one pair in g. Asterisk indicates significance from 0: *p<0.05; **p<0.01.

**Fig. S8.**
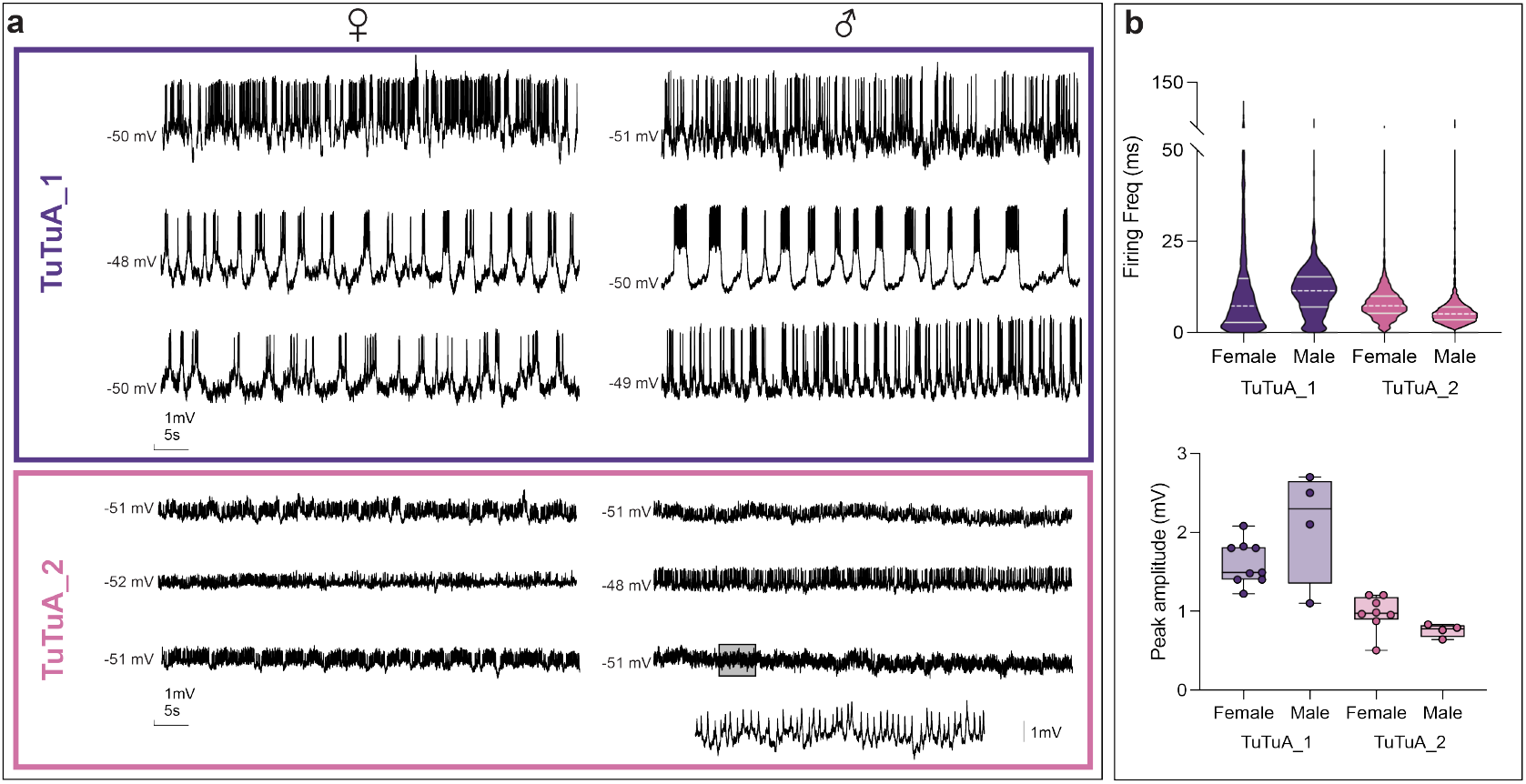
Recordings from TuTuA_1 and TuTuA_2 in males and females. Each trace is one-minute recording from one cell. TuTuA_1 displayed the larger action potential amplitude compared to TuTuA_2, with similar properties between males and females. Inset recording from a TuTuA_2 neuron is from the highlighted region of the last male recording. (b) Analysis of the firing frequency and peak amplitude of TuTuA_1 and TuTuA_2 recordings in males and females. The instantaneous action potential frequency was calculated for about one minute in each cell (TuTuA_1: Female, n = 1536, Male, n = 1965; TuTuA_2: Female, n = 1198, Male, n = 1185). The action potential amplitude was averaged from 20-30 individual events in each cell (each dot represents 1 cell), and measured as the difference between the threshold and peak (TuTuA_1: Female, n = 9, Male, n = 4; TuTuA_2: Female, n = 8, Male, n = 4). The firing frequency was more variable in the TuTuA_1 recordings than in the TuTuA_2 recordings in both males and females. Additionally, the amplitude from TuTuA_1 was larger compared to TuTuA_2 in both males and females. However, the action potential is dramatically slower in male TuTuA_2 neurons. Box-and-whisker and violin plots show median and IQR; whiskers or ends of the violin show range.

**Fig. S9.**
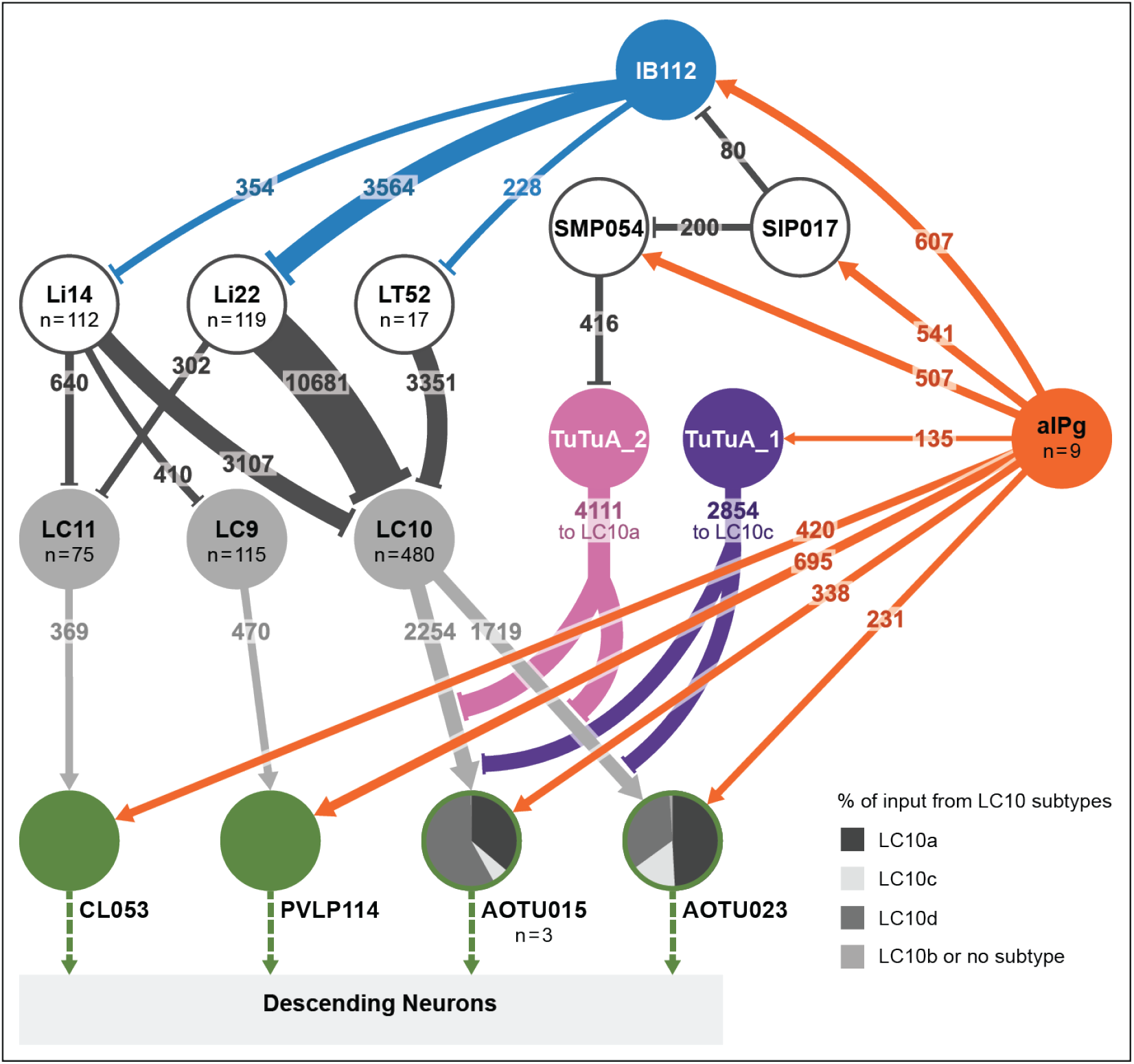
Circuit diagram of aIPg modulation of visual processing. A detailed circuit map of the mechanisms detailed in Figure 1a and Figure 6. This diagram shows additional downstream targets of aIPg including those involved in regulating information flow from LC9 and LC11. The diagram also illustrates that CL053, PVLP114, AOTU015 and AOTU023 are each upstream of descending interneurons (DNs) that traverse the neck into the ventral nerve cord where they likely regulate motor action. Each of these neurons connect to largely non-overlapping sets of DNs, implying that these parallel pathways control different motor actions. Numbers within arrows indicate synapses numbers. The top six downstream targets of aIPg are represented in this diagram: (1) PVLP114; (2) IB112; (3) SMP054; (4) SIP017; (5) CL053; and (6) AOTU015. Li22 devotes 66% (10,627/16,168) of its synapses going to any LC cell type to LC10 and provides input to all LC10 subtypes; on average the number of Li22 inputs per cell to each LC10 subtype are as follows: LC10a, 23; LC10b, 12; LC10c, 27; and LC10d, 32. Li14 distributes its output more broadly with only 20% of its output to LC neurons going to LC10 with a more biased distribution between subtypes than Li22; on average the number of Li14 inputs per cell to each LC10 subtype are as follows: LC10a, 14; LC10b, 25; LC10c, 2; and LC10d, 5. LT52 devotes 57% of its synapses in the lobula that go to LC neurons to LC10 subtypes with a strong bias to LC10b and LC10d; on average the number of LT52 inputs per cell to each LC10 subtype are as follows: LC10a, 5; LC10b, 27; LC10c, <1; and LC10d, 12.

## Bibliography

1. John Morgan Allman. Evolving brains. Number 68 in Scientific American library series. Scientifc American Library, New York, 2000. ISBN 978-0-7167-5076-5.

2. Timothy A Currier, Michelle M Pang, and Thomas R Clandinin. Visual processing in the fly, from photoreceptors to behavior. Genetics, 224(2):iyad064, June 2023. ISSN 1943-2631. doi:10.1093/genetics/iyad064.

3. Kit D. Longden, Edward M. Rogers, Aljoscha Nern, Heather Dionne, and Michael B. Reiser. Different spectral sensitivities of ON-and OFF-motion pathways enhance the detection of approaching color objects in Drosophila. Nature Communications, 14(1):7693, November 2023. ISSN 2041-1723. doi:10.1038/s41467-023-43566-8.

4. Melvyn A. Goodale and A. David Milner. Separate visual pathways for perception and action. Trends in Neurosciences, 15(1):20–25, January 1992. ISSN 0166-2236. doi:10.1016/0166-2236(92)90344-8.

5. Bevil R. Conway. Color signals through dorsal and ventral visual pathways. Visual Neuroscience, 31(2):197–209, March 2014. ISSN 0952-5238, 1469-8714. doi:10.1017/S0952523813000382.

6. Kazunori Shinomiya, Aljoscha Nern, Ian A. Meinertzhagen, Stephen M. Plaza, and Michael B. Reiser. Neuronal circuits integrating visual motion information in Drosophila melanogaster. Current Biology, 32(16):3529–3544.e2, August 2022. ISSN 0960-9822. doi:10.1016/j.cub.2022.06.061.

7. János A. Perge, Bart G. Borghuis, Roger J. E. Bours, Martin J. M. Lankheet, and Richard J. A. van Wezel. Temporal dynamics of direction tuning in motion-sensitive macaque area MT. Journal of Neurophysiology, 93(4):2104–2116, April 2005. ISSN 0022-3077. doi:10.1152/jn.00601.2004.

8. Dario L. Ringach, Michael J. Hawken, and Robert Shapley. Dynamics of orientation tuning in macaque primary visual cortex. Nature, 387(6630):281–284, May 1997. ISSN 1476-4687. doi:10.1038/387281a0.

9. Dennis M. Dacey. Parallel pathways for spectral coding in primate retina. Annual Review of Neuroscience, 23(1):743–775, 2000. doi:10.1146/annurev.neuro.23.1.743.

10. Maximilian Joesch and Markus Meister. A neuronal circuit for colour vision based on rod–cone opponency. Nature, 532(7598):236–239, April 2016. ISSN 1476-4687. doi:10.1038/nature17158.

11. Fabrizio Gabbiani, Holger G. Krapp, and Gilles Laurent. Computation of object approach by a wide-field, motion-sensitive neuron. Journal of Neuroscience, 19(3):1122–1141, February 1999. ISSN 0270-6474, 1529-2401. doi:10.1523/JNEUROSCI.19-03-01122.1999.

12. Pierre-Olivier Polack, Jonathan Friedman, and Peyman Golshani. Cellular mechanisms of brain state–dependent gain modulation in visual cortex. Nature Neuroscience, 16(9):1331– 1339, September 2013. ISSN 1546-1726. doi:10.1038/nn.3464.

13. Carsen Stringer, Michalis Michaelos, Dmitri Tsyboulski, Sarah E. Lindo, and Marius Pachitariu. High-precision coding in visual cortex. Cell, 184(10):2767–2778.e15, May 2021. ISSN 00928674. doi:10.1016/j.cell.2021.03.042.

14. Lilach Avitan and Carsen Stringer. Not so spontaneous: Multi-dimensional representations of behaviors and context in sensory areas. Neuron, 110(19):3064–3075, October 2022. ISSN 0896-6273. doi:10.1016/j.neuron.2022.06.019.

15. J. L. Herrero, M. J. Roberts, L. S. Delicato, M. A. Gieselmann, P. Dayan, and A. Thiele. Acetylcholine contributes through muscarinic receptors to attentional modulation in V1. Nature, 454(7208):1110–1114, August 2008. ISSN 1476-4687. doi:10.1038/nature07141.

16. Michael Goard and Yang Dan. Basal forebrain activation enhances cortical coding of natural scenes. Nature Neuroscience, 12(11):1444–1449, November 2009. ISSN 1546-1726. doi:10.1038/nn.2402.

17. Cristopher M. Niell and Michael P. Stryker. Modulation of visual responses by behavioral state in mouse visual cortex. Neuron, 65(4):472–479, February 2010. ISSN 08966273. doi:10.1016/j.neuron.2010.01.033.

18. Ming Wu, Aljoscha Nern, W Ryan Williamson, Mai M Morimoto, Michael B Reiser, Gwyneth M Card, and Gerald M Rubin. Visual projection neurons in the Drosophila lobula link feature detection to distinct behavioral programs. eLife, 5:e21022, December 2016. ISSN 2050-084X. doi:10.7554/eLife.21022.

19. Mai M Morimoto, Aljoscha Nern, Arthur Zhao, Edward M Rogers, Allan M Wong, Mathew D Isaacson, Davi D Bock, Gerald M Rubin, and Michael B Reiser. Spatial readout of visual looming in the central brain of Drosophila. eLife, 9:e57685, November 2020. ISSN 2050-084X. doi:10.7554/eLife.57685.

20. Jan M. Ache, Jason Polsky, Shada Alghailani, Ruchi Parekh, Patrick Breads, Martin Y. Peek, Davi D. Bock, Catherine R. Von Reyn, and Gwyneth M. Card. Neural basis for looming size and velocity encoding in the Drosophila giant fiber escape pathway. Current Biology, 29(6): 1073–1081.e4, March 2019. ISSN 09609822. doi:10.1016/j.cub.2019.01.079.

21. Hideo Otsuna and Kei Ito. Systematic analysis of the visual projection neurons of Drosophila melanogaster. I. Lobula-specific pathways. The Journal of Comparative Neurology, 497(6): 928–958, August 2006. ISSN 0021-9967. doi:10.1002/cne.21015.

22. Mehmet F. Keleş and Mark A. Frye. Object-detecting neurons in Drosophila. Current biology: CB, 27(5):680–687, March 2017. ISSN 1879-0445. doi:10.1016/j.cub.2017.01.012.

23. Nathan C. Klapoetke, Aljoscha Nern, Edward M. Rogers, Gerald M. Rubin, Michael B. Reiser, and Gwyneth M. Card. A functionally ordered visual feature map in the Drosophila brain. Neuron, 110(10):1700–1711.e6, May 2022. ISSN 08966273. doi:10.1016/j.neuron.2022.02.013.

24. Louis K Scheffer, C Shan Xu, Michal Januszewski, Zhiyuan Lu, Shin-ya Takemura, Kenneth J Hayworth, Gary B Huang, Kazunori Shinomiya, Jeremy Maitlin-Shepard, Stuart Berg, Jody Clements, Philip M Hubbard, William T Katz, Lowell Umayam, Ting Zhao, David Ackerman, Tim Blakely, John Bogovic, Tom Dolafi, Dagmar Kainmueller, Takashi Kawase, Khaled A Khairy, Laramie Leavitt, Peter H Li, Larry Lindsey, Nicole Neubarth, Donald J Olbris, Hideo Otsuna, Eric T Trautman, Masayoshi Ito, Alexander S Bates, Jens Goldammer, Tanya Wolff, Robert Svirskas, Philipp Schlegel, Erika Neace, Christopher J Knecht, Chelsea X Alvarado, Dennis A Bailey, Samantha Ballinger, Jolanta A Borycz, Brandon S Canino, Natasha Cheatham, Michael Cook, Marisa Dreher, Octave Duclos, Bryon Eubanks, Kelli Fairbanks, Samantha Finley, Nora Forknall, Audrey Francis, Gary Patrick Hopkins, Emily M Joyce, SungJin Kim, Nicole A Kirk, Julie Kovalyak, Shirley A Lauchie, Alanna Lohff, Charli Maldonado, Emily A Manley, Sari McLin, Caroline Mooney, Miatta Ndama, Omotara Ogundeyi, Nneoma Okeoma, Christopher Ordish, Nicholas Padilla, Christopher M Patrick, Tyler Paterson, Elliott E Phillips, Emily M Phillips, Neha Rampally, Caitlin Ribeiro, Madelaine K Robertson, Jon Thomson Rymer, Sean M Ryan, Megan Sammons, Anne K Scott, Ashley L Scott, Aya Shinomiya, Claire Smith, Kelsey Smith, Natalie L Smith, Margaret A Sobeski, Alia Suleiman, Jackie Swift, Satoko Takemura, Iris Talebi, Dorota Tarnogorska, Emily Tenshaw, Temour Tokhi, John J Walsh, Tansy Yang, Jane Anne Horne, Feng Li, Ruchi Parekh, Patricia K Rivlin, Vivek Jayaraman, Marta Costa, Gregory SXE Jefferis, Kei Ito, Stephan Saalfeld, Reed George, Ian A Meinertzhagen, Gerald M Rubin, Harald F Hess, Viren Jain, and Stephen M Plaza. A connectome and analysis of the adult Drosophila central brain. eLife, 9:e57443, September 2020. ISSN 2050-084X. doi:10.7554/eLife.57443.

25. Gaby Maimon, Andrew D. Straw, and Michael H. Dickinson. Active flight increases the gain of visual motion processing in Drosophila. Nature Neuroscience, 13(3):393–399, March 2010. ISSN 1546-1726. doi:10.1038/nn.2492.

26. R. Rosner, M. Egelhaaf, and A.-K. Warzecha. Behavioural state affects motion-sensitive neurones in the fly visual system. Journal of Experimental Biology, 213(2):331–338, January 2010. ISSN 0022-0949. doi:10.1242/jeb.035386.

27. Kit D. Longden and Holger G. Krapp. State-dependent performance of optic-flow processing interneurons. Journal of Neurophysiology, 102(6):3606–3618, December 2009. ISSN 0022-3077. doi:10.1152/jn.00395.2009.

28. Kit Longden and Holger Krapp. Octopaminergic modulation of temporal frequency coding in an identified optic flow-processing interneuron. Frontiers in Systems Neuroscience, 4, 2010. ISSN 1662-5137.

29. James A. Strother, Shiuan-Tze Wu, Edward M. Rogers, Jessica L. M. Eliason, Allan M. Wong, Aljoscha Nern, and Michael B. Reiser. Behavioral state modulates the ON visual motion pathway of Drosophila. Proceedings of the National Academy of Sciences, 115(1): E102–E111, January 2018. doi:10.1073/pnas.1703090115.

30. Jan M. Ache, Shigehiro Namiki, Allen Lee, Kristin Branson, and Gwyneth M. Card. State-dependent decoupling of sensory and motor circuits underlies behavioral flexibility in Drosophila. Nature Neuroscience, 22(7):1132–1139, July 2019. ISSN 1546-1726. doi:10.1038/s41593-019-0413-4.

31. Tess B. Oram and Gwyneth M. Card. Context-dependent control of behavior in Drosophila. Current Opinion in Neurobiology, 73:102523, April 2022. ISSN 0959-4388. doi:10.1016/j.conb.2022.02.003.

32. Sweta Agrawal, Steve Safarik, and Michael Dickinson. The relative roles of vision and chemosensation in mate recognition of Drosophila melanogaster. Journal of Experimental Biology, 217(15):2796–2805, August 2014. ISSN 0022-0949. doi:10.1242/jeb.105817.

33. Sweta Agrawal and Michael H. Dickinson. The effects of target contrast on Drosophila courtship. Journal of Experimental Biology, 222(16):jeb203414, August 2019. ISSN 0022-0949. doi:10.1242/jeb.203414.

34. Susanne C. Hoyer, Andreas Eckart, Anthony Herrel, Troy Zars, Susanne A. Fischer, Shannon L. Hardie, and Martin Heisenberg. Octopamine in male aggression of Drosophila. Current Biology, 33(18):3896–3910.e7, September 2023. ISSN 0960-9822. doi:10.1016/j.cub.2023.08.034.

35. Eric D Hoopfer. Neural control of aggression in Drosophila. Current Opinion in Neurobiology, 38:109–118, June 2016. ISSN 0959-4388. doi:10.1016/j.conb.2016.04.007.

36. Ryosuke Tanaka and Damon A. Clark. Object-displacement-sensitive visual neurons drive freezing in Drosophila. Current Biology, 30(13):2532–2550.e8, July 2020. ISSN 0960-9822. doi:10.1016/j.cub.2020.04.068.

37. Mehmet F. Keleş, Ben J. Hardcastle, Carola Städele, Qi Xiao, and Mark A. Frye. Inhibitory interactions and columnar inputs to an object motion detector in Drosophila. Cell Reports, 30(7):2115–2124.e5, February 2020. ISSN 2211-1247. doi:10.1016/j.celrep.2020.01.061.

38. Salil S. Bidaye, Meghan Laturney, Amy K. Chang, Yuejiang Liu, Till Bockemuhl, Ansgar Buschges, and Kristin Scott. Two brain pathways initiate distinct forward walking programs in Drosophila. Neuron, 108(3):469–485.e8, November 2020. ISSN 0896-6273. doi:10.1016/j.neuron.2020.07.032.

39. Soh Kohatsu and Daisuke Yamamoto. Visually induced initiation of Drosophila innate courtship-like following pursuit is mediated by central excitatory state. Nature Communications, 6(1):6457, March 2015. ISSN 2041-1723. doi:10.1038/ncomms7457.

40. Inês M.A. Ribeiro, Michael Drews, Armin Bahl, Christian Machacek, Alexander Borst, and Barry J. Dickson. Visual projection neurons mediating directed courtship in Drosophila. Cell, 174(3):607–621.e18, July 2018. ISSN 00928674. doi:10.1016/j.cell.2018.06.020.

41. Tom Hindmarsh Sten, Rufei Li, Adriane Otopalik, and Vanessa Ruta. Sexual arousal gates visual processing during Drosophila courtship. Nature, 595(7868):549–553, July 2021. ISSN 1476-4687. doi:10.1038/s41586-021-03714-w.

42. Carola Städele, Mehmet F. Keleş, Jean-Michel Mongeau, and Mark A. Frye. Non-canonical receptive field properties and neuromodulation of feature-detecting neurons in flies. Current Biology, 30(13):2508–2519.e6, July 2020. ISSN 0960-9822. doi:10.1016/j.cub.2020.04.069.

43. Eric D Hoopfer, Yonil Jung, Hidehiko K Inagaki, Gerald M Rubin, and David J Anderson. P1 interneurons promote a persistent internal state that enhances inter-male aggression in Drosophila. eLife, 4:e11346, December 2015. ISSN 2050-084X. doi:10.7554/eLife.11346.

44. Sebastian Cachero, Aaron D. Ostrovsky, Jai Y. Yu, Barry J. Dickson, and Gregory S. X. E. Jefferis. Sexual dimorphism in the fly brain. Current Biology, 20(18):1589–1601, September 2010. ISSN 0960-9822. doi:10.1016/j.cub.2010.07.045.

45. Jai Y. Yu, Makoto I. Kanai, Ebru Demir, Gregory S. X. E. Jefferis, and Barry J. Dickson. Cellular organization of the neural circuit that drives Drosophila courtship behavior. Current Biology, 20(18):1602–1614, September 2010. ISSN 0960-9822. doi:10.1016/j.cub.2010.08.025.

46. Ken-ichi Kimura, Tomoaki Hachiya, Masayuki Koganezawa, Tatsunori Tazawa, and Daisuke Yamamoto. Fruitless and doublesex coordinate to generate male-specific neurons that can initiate courtship. Neuron, 59(5):759–769, September 2008. ISSN 0896-6273. doi:10.1016/j.neuron.2008.06.007.

47. Catherine E Schretter, Yoshinori Aso, Alice A Robie, Marisa Dreher, Michael-John Dolan, Nan Chen, Masayoshi Ito, Tansy Yang, Ruchi Parekh, Kristin M Branson, and Gerald M Rubin. Cell types and neuronal circuitry underlying female aggression in Drosophila. eLife, 9:e58942, November 2020. ISSN 2050-084X. doi:10.7554/eLife.58942.

48. David Deutsch, Diego Pacheco, Lucas Encarnacion-Rivera, Talmo Pereira, Ramie Fathy, Jan Clemens, Cyrille Girardin, Adam Calhoun, Elise Ireland, Austin Burke, Sven Dorkenwald, Claire McKellar, Thomas Macrina, Ran Lu, Kisuk Lee, Nico Kemnitz, Dodam Ih, Manuel Castro, Akhilesh Halageri, Chris Jordan, William Silversmith, Jingpeng Wu, H Sebastian Seung, and Mala Murthy. The neural basis for a persistent internal state in Drosophila females. eLife, 9:e59502, November 2020. ISSN 2050-084X. doi:10.7554/eLife.59502.

49. Hui Chiu, Alice A. Robie, Kristin M. Branson, Tanvi Vippa, Samantha Epstein, Gerald M. Rubin, David J. Anderson, and Catherine E. Schretter. Cell type-specific contributions to a persistent aggressive internal state in female Drosophila. eLife, 12, December 2023. doi:10.7554/eLife.88598.2.

50. B. T. Bloomquist, R. D. Shortridge, S. Schneuwly, M. Perdew, C. Montell, H. Steller, G. Rubin, and W. L. Pak. Isolation of a putative phospholipase c gene of drosophila, norpA, and its role in phototransduction. Cell, 54(5):723–733, August 1988. ISSN 0092-8674. doi:10.1016/S0092-8674(88)80017-5.

51. Helen H. Yang, Luke E. Brezovec, Laia Serratosa Capdevila, Quinn X. Vanderbeck, Atsuko Adachi, Richard S. Mann, and Rachel I. Wilson. Fine-grained descending control of steering in walking Drosophila, October 2023.

52. Aleksandr Rayshubskiy, Stephen L. Holtz, Isabel D’Alessandro, Anna A. Li, Quinn X. Vanderbeck, Isabel S. Haber, Peter W. Gibb, and Rachel I. Wilson. Neural circuit mechanisms for steering control in walking Drosophila, July 2020.

53. Wendy W. Liu and Rachel I. Wilson. Glutamate is an inhibitory neurotransmitter in the Drosophila olfactory system. Proceedings of the National Academy of Sciences, 110(25): 10294–10299, June 2013. doi:10.1073/pnas.1220560110.

54. Aljoscha Nern et al. Connectome-driven neural inventory of a complete visual system, 2024. in prep.

55. Sven Dorkenwald, Arie Matsliah, Amy R. Sterling, Philipp Schlegel, Szi-chieh Yu, Claire E. McKellar, Albert Lin, Marta Costa, Katharina Eichler, Yijie Yin, Will Silversmith, Casey Schneider-Mizell, Chris S. Jordan, Derrick Brittain, Akhilesh Halageri, Kai Kuehner, Oluwaseun Ogedengbe, Ryan Morey, Jay Gager, Krzysztof Kruk, Eric Perlman, Runzhe Yang, David Deutsch, Doug Bland, Marissa Sorek, Ran Lu, Thomas Macrina, Kisuk Lee, J. Alexander Bae, Shang Mu, Barak Nehoran, Eric Mitchell, Sergiy Popovych, Jingpeng Wu, Zhen Jia, Manuel Castro, Nico Kemnitz, Dodam Ih, Alexander Shakeel Bates, Nils Eckstein, Jan Funke, Forrest Collman, Davi D. Bock, Gregory S. X. E. Jefferis, H. Sebastian Seung, Mala Murthy, and the FlyWire Consortium. Neuronal wiring diagram of an adult brain, July 2023.

56. Marie P. Suver, Akira Mamiya, and Michael H. Dickinson. Octopamine neurons mediate flight-induced modulation of visual processing in Drosophila. Current Biology, 22(24):2294– 2302, December 2012. ISSN 0960-9822. doi:10.1016/j.cub.2012.10.034.

57. Yuta Mabuchi, Xinyue Cui, Lily Xie, Haein Kim, Tianxing Jiang, and Nilay Yapici. Visual feedback neurons fine-tune Drosophila male courtship via GABA-mediated inhibition. Current Biology, 33(18):3896–3910.e7, September 2023. ISSN 0960-9822. doi:10.1016/j.cub.2023.08.034.

58. Oliver Braganza and Heinz Beck. The circuit motif as a conceptual tool for multilevel neuroscience. Trends in Neurosciences, 41(3):128–136, March 2018. ISSN 0166-2236. doi:10.1016/j.tins.2018.01.002.

59. Andrew C. Lin, Alexei M. Bygrave, Alix de Calignon, Tzumin Lee, and Gero Miesenböck. Sparse, decorrelated odor coding in the mushroom body enhances learned odor discrimination. Nature Neuroscience, 17(4):559–568, April 2014. ISSN 1546-1726. doi:10.1038/nn.3660.

60. Zeno Jonke, Robert Legenstein, Stefan Habenschuss, and Wolfgang Maass. Feedback inhibition shapes emergent computational properties of cortical microcircuit motifs. Journal of Neuroscience, 37(35):8511–8523, August 2017. ISSN 0270-6474, 1529-2401. doi:10.1523/JNEUROSCI.2078-16.2017.

61. Heather Dionne, Karen L Hibbard, Amanda Cavallaro, Jui-Chun Kao, and Gerald M Rubin. Genetic reagents for making split-GAL4 lines in Drosophila. Genetics, 209(1):31–35, May 2018. ISSN 1943-2631. doi:10.1534/genetics.118.300682.

62. Alice A. Robie et al. The fly disco: Instrument and analysis pipeline for optogenetic manipulation and fine-grained analysis of fly behavior, 2024. in prep.

63. Mayank Kabra, Alice A. Robie, Marta Rivera-Alba, Steven Branson, and Kristin Branson. JAABA: interactive machine learning for automatic annotation of animal behavior. Nature Methods, 10(1):64–67, January 2013. ISSN 1548-7105. doi:10.1038/nmeth.2281.

64. Alice A. Robie, Jonathan Hirokawa, Austin W. Edwards, Lowell A. Umayam, Allen Lee, Mary L. Phillips, Gwyneth M. Card, Wyatt Korff, Gerald M. Rubin, Julie H. Simpson, Michael B. Reiser, and Kristin Branson. Mapping the neural substrates of behavior. Cell, 170(2):393–406.e28, July 2017. ISSN 0092-8674. doi:10.1016/j.cell.2017.06.032.

65. James A. Strother, Aljoscha Nern, and Michael B. Reiser. Direct observation of ON and OFF pathways in the Drosophila visual system. Current Biology, 24(9):976–983, May 2014. ISSN 0960-9822. doi:10.1016/j.cub.2014.03.017.

66. Richard J. D. Moore, Gavin J. Taylor, Angelique C. Paulk, Thomas Pearson, Bruno van Swinderen, and Mandyam V. Srinivasan. FicTrac: A visual method for tracking spherical motion and generating fictive animal paths. Journal of Neuroscience Methods, 225:106– 119, March 2014. ISSN 0165-0270. doi:10.1016/j.jneumeth.2014.01.010.

67. Dmitriy Aronov and David W. Tank. Engagement of neural circuits underlying 2D spatial navigation in a rodent virtual reality system. Neuron, 84(2):442–456, October 2014. ISSN 0896-6273. doi:10.1016/j.neuron.2014.08.042.

68. Yoshinori Aso, Robert P Ray, Xi Long, Daniel Bushey, Karol Cichewicz, Teri-TB Ngo, Brandi Sharp, Christina Christoforou, Amy Hu, Andrew L Lemire, Paul Tillberg, Jay Hirsh, Ashok Litwin-Kumar, and Gerald M Rubin. Nitric oxide acts as a cotransmitter in a subset of dopaminergic neurons to diversify memory dynamics. eLife, 8:e49257, November 2019. ISSN 2050-084X. doi:10.7554/eLife.49257.

69. Yoshinori Aso, Daisuke Hattori, Yang Yu, Rebecca M Johnston, Nirmala A Iyer, Teri-TB Ngo, Heather Dionne, LF Abbott, Richard Axel, Hiromu Tanimoto, and Gerald M Rubin. The neuronal architecture of the mushroom body provides a logic for associative learning. eLife, 3:e04577, December 2014. ISSN 2050-084X. doi:10.7554/eLife.04577.

70. Aljoscha Nern, Barret D. Pfeiffer, and Gerald M. Rubin. Optimized tools for multicolor stochastic labeling reveal diverse stereotyped cell arrangements in the fly visual system. Proceedings of the National Academy of Sciences, 112(22):E2967–E2976, June 2015. doi:10.1073/pnas.1506763112.

71. Gabriella R Sterne, Hideo Otsuna, Barry J Dickson, and Kristin Scott. Classification and genetic targeting of cell types in the primary taste and premotor center of the adult Drosophila brain. eLife, 10:e71679, September 2021. ISSN 2050-084X. doi:10.7554/eLife.71679.

72. Geoffrey W Meissner, Aljoscha Nern, Zachary Dorman, Gina M DePasquale, Kaitlyn Forster, Theresa Gibney, Joanna H Hausenfluck, Yisheng He, Nirmala A Iyer, Jennifer Jeter, Lauren Johnson, Rebecca M Johnston, Kelley Lee, Brian Melton, Brianna Yarbrough, Christopher T Zugates, Jody Clements, Cristian Goina, Hideo Otsuna, Konrad Rokicki, Robert R Svirskas, Yoshinori Aso, Gwyneth M Card, Barry J Dickson, Erica Ehrhardt, Jens Goldammer, Masayoshi Ito, Dagmar Kainmueller, Wyatt Korff, Lisa Mais, Ryo Minegishi, Shigehiro Namiki, Gerald M Rubin, Gabriella R Sterne, Tanya Wolff, Oz Malkesman, and FlyLight Project Team. A searchable image resource of Drosophila GAL4 driver expression patterns with single neuron resolution. eLife, 12:e80660, February 2023. ISSN 2050-084X. doi:10.7554/eLife.80660.

73. Geoffrey W. Meissner, Allison Vannan, Jennifer Jeter, Megan Atkins, Shelby Bowers, Kari Close, Gina M. DePasquale, Zachary Dorman, Kaitlyn Forster, Jaye Anne Beringer, Theresa V. Gibney, Asish Gulati, Joanna H. Hausenfluck, Yisheng He, Kristin Henderson, Lauren Johnson, Rebecca M. Johnston, Gudrun Ihrke, Nirmala Iyer, Rachel Lazarus, Kelley Lee, Hsing-Hsi Li, Hua-Peng Liaw, Brian Melton, Scott Miller, Reeham Motaher, Alexandra Novak, Omatara Ogundeyi, Alyson Petruncio, Jacquelyn Price, Sophia Protopapas, Susana Tae, Athreya Tata, Jennifer Taylor, Rebecca Vorimo, Brianna Yarbrough, Kevin Xiankun Zeng, Christopher T. Zugates, Heather Dionne, Claire Angstadt, Kelly Ashley, Amanda Cavallaro, Tam Dang, Guillermo A. Gonzalez, Karen L. Hibbard, Cuizhen Huang, Jui-Chun Kao, Todd Laverty, Monti Mercer, Brenda Perez, Scarlett Pitts, Danielle Ruiz, Viruthika Vallanadu, Grace Zhiyu Zheng, Cristian Goina, Hideo Otsuna, Konrad Rokicki, Robert R. Svirskas, Han SJ Cheong, Michael-John Dolan, Erica Ehrhardt, Kai Feng, Basel El Galfi, Jens Goldammer, Nan Hu, Masayoshi Ito, Claire McKellar, Ryo Minegishi, Shigehiro Namiki, Aljoscha Nern, Catherine E. Schretter, Gabriella R. Sterne, Lalanti Venkatasubramanian, Kaiyu Wang, Tanya Wolff, Ming Wu, Reed George, Oz Malkesman, Yoshinori Aso, Gwyneth M. Card, Barry J. Dickson, Wyatt Korff, Kei Ito, James W. Truman, Marta Zlatic, Gerald M. Rubin, and FlyLight Project Team. A split-GAL4 driver line resource for Drosophila CNS cell types, January 2024.

74. Zhihao Zheng, J. Scott Lauritzen, Eric Perlman, Camenzind G. Robinson, Matthew Nichols, Daniel Milkie, Omar Torrens, John Price, Corey B. Fisher, Nadiya Sharifi, Steven A. Calle-Schuler, Lucia Kmecova, Iqbal J. Ali, Bill Karsh, Eric T. Trautman, John A. Bogovic, Philipp Hanslovsky, Gregory S. X. E. Jefferis, Michael Kazhdan, Khaled Khairy, Stephan Saalfeld, Richard D. Fetter, and Davi D. Bock. A complete electron microscopy volume of the brain of adult Drosophila melanogaster. Cell, 174(3):730–743.e22, July 2018. ISSN 0092-8674, 1097-4172. doi:10.1016/j.cell.2018.06.019.

75. Barret D Pfeiffer, Teri-T B Ngo, Karen L Hibbard, Christine Murphy, Arnim Jenett, James W Truman, and Gerald M Rubin. Refinement of tools for targeted gene expression in Drosophila. Genetics, 186(2):735–755, October 2010. ISSN 1943-2631. doi:10.1534/genetics.110.119917.

76. John A. Bogovic, Hideo Otsuna, Larissa Heinrich, Masayoshi Ito, Jennifer Jeter, Geoffrey Meissner, Aljoscha Nern, Jennifer Colonell, Oz Malkesman, Kei Ito, and Stephan Saalfeld. An unbiased template of the Drosophila brain and ventral nerve cord. PloS One, 15(12): e0236495, 2020. ISSN 1932-6203. doi:10.1371/journal.pone.0236495.

